# Environmental flexibility does not explain metabolic robustness

**DOI:** 10.1101/2020.10.04.325407

**Authors:** Julian Libiseller-Egger, Ben Coltman, Matthias P. Gerstl, Jürgen Zanghellini

**Affiliations:** Austrian Centre of Industrial Biotechnology, A-1190 Vienna, Austria; University of Natural Resources and Life Sciences, A-1190 Vienna, Austria; Department of Biotechnology, University of Natural Resources and Life Sciences, A-1190 Vienna, Austria; Department of Analytical Chemistry, University of Vienna, A-1090 Vienna, Austria

## Abstract

Cells show remarkable resilience against genetic and environmental perturbations. However, its evolutionary origin remains obscure. In order to leverage methods of systems biology for examining cellular robustness, a computationally accessible way of quantification is needed. Here, we present an unbiased metric of structural robustness in genome-scale metabolic models based on concepts prevalent in reliability engineering and fault analysis.

The probability of failure (PoF) is defined as the (weighted) portion of all possible combinations of loss-of-function mutations that disable network functionality. It can be exactly determined, if all essential reactions, synthetic lethal pairs of reactions, synthetic lethal triplets of reactions etc., are known. In theory, these minimal cut sets (MCSs) can be calculated for any network, but for large models the problem remains computationally intractable. Herein, we demonstrate that even at the genome scale only the lowest-cardinality MCSs are required to efficiently approximate the PoF with reasonable accuracy.

We analysed the robustness of 489 *E. coli, Shigella, Salmonella*, and fungal genome-scale metabolic models (GSMMs). In contrast to the popular “congruence theory”, which explains the origin of genetic robustness as a byproduct of selection for environmental flexibility, we found no correlation between network robustness and the diversity of growth-supporting environments. On the contrary, our analysis indicates that amino acid synthesis rather than carbon metabolism dominates metabolic robustness.

## INTRODUCTION

“Robustness” is a system’s intrinsic ability to maintain functionality under perturbation. Due to the generality of this definition, a consensus on an exact quantitative metric, especially in a biological context, has yet to emerge [1–4]. Aside from biology, robustness and reliability are key design goals in many fields of engineering and particularly paramount in the development of safety-critical systems [5]. Thus, some of the theoretical groundwork relevant in these areas can be borrowed for a systems biology approach [2, 3].

### Box 1

**List of symbols**

*m*: Number of minimal cut sets (MCSs) disabling growth in a metabolic network
*m*_*i*_: Number of MCSs with cardinalities up to *i*
*d*_*m*_: Cardinality up to which all MCSs have been calculated (i.e. for *d*_*m*_ = 3 all *m*_3_ MCSs with up to three reactions are known)
*d*_0_: Cardinality up to which all supersets of all known MCSs are considered by `pof2.0`
*f*_*d*_: Failure frequency at *d* random reaction deletions (i.e. the chance of a metabolic network not being able to grow given loss-of-function (LOF) mutations in *d* random reactions)
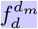: Approximated failure frequency at *d* random reaction deletions computed with all MCSs up to cardinality *d*_*m*_
*F*: Probability of failure (PoF) (probability of a metabolic network acquiring a lethal combination of LOF mutations)
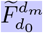: PoF estimate calculated by `pof2.0` given all MCSs up to cardinality *d*_*m*_ and considering all possible unions of MCSs up to cardinality *d*_0_

Common basic reliability measures consider only the interaction of binary-state components, i.e. components which are either fully functional or fully dysfunctional. In such an analysis, a network is deemed functional if there exists at least one path from the input node(s) to the output node(s). Some early efforts to quantify structural robustness in metabolic networks follow the same principle. They correlate robustness with the number of unique, minimal pathways [6], that provide a given function (in the context of metabolism usually cell growth) [7–9].

Consistent with the characterisation of “functionality” as cell growth, “dysfunctionality” (i.e. the lack of growth) is caused by lethal combinations of reaction knockouts, which could arise via loss-of-function (LOF) mutations disabling vital enzymatic activity. Similarly to how all paths leading from input nodes to output nodes can be determined computationally [10], the same can be done for all ways that block these paths (i.e. lethal combinations of reaction deletions) [11]. Analogously, knowledge of these knockout combinations, termed minimal cut sets (MCSs) [12], allows for computing the network’s probability of failure (PoF), which – in mathematical terms – is the complement of robustness [13].

As we have shown previously, the great advantage in using the PoF as a surrogate for robustness lies in the fact that only the lowest cardinality-MCS are sufficient to approximate it with negligible error [13]. As efficient algorithms for finding low-cardinality MCSs are available [14, 15], this allows for the examination of robustness in genome-scale metabolic models (GSMMs).

### Box 2

**Probability of failure (PoF) by example**

The toy network composed of five irreversible and one reversible reaction depicted in **Fig. 1a** produces biomass via *r*_6_. Any combination of LOF mutations encompassing the essential reaction *r*_6_, one of the synthetic lethal pairs {*r*_1_, *r*_2_}, {*r*_4_, *r*_5_}, or one of the synthetic lethal triplets {*r*_1_, *r*_3_, *r*_5_}, {*r*_2_, *r*_3_, *r*_4_} is lethal. The failure frequency, *f*_*d*_ (i.e. the probability that a given number of *d* LOF mutations results in cell death), is illustrated in **Fig. 1b**. In general, the failure frequency for any fixed *d* is dominated by single and double LOF mutations (see the hatched bars in **Fig. 1b**), while the contribution of higher order mutations is relatively small. For instance, in **Fig. 1b** all *f*_*d*_ – except for *f*_3_ – can be computed exactly with the knowledge of the single and double LOF mutations alone as all *f*_*d*_ with *d >* 3 are already determined to be 1 by their respective contributions.

For a constant mutation rate *p* = 0.1 the likelihood that one, two, or more LOF mutations occur is binomially distributed, and the resulting expected value of the probability of a lethal set of mutations (i.e. the PoF) is *p* + 2*p*^2^ – 7*p*^4^ + 7*p*^5^ – 2*p*^6^ = 0.11936 (see **Supplementary Table 2** for details). Note that it is independent of the network’s size.

**Fig. 1.**
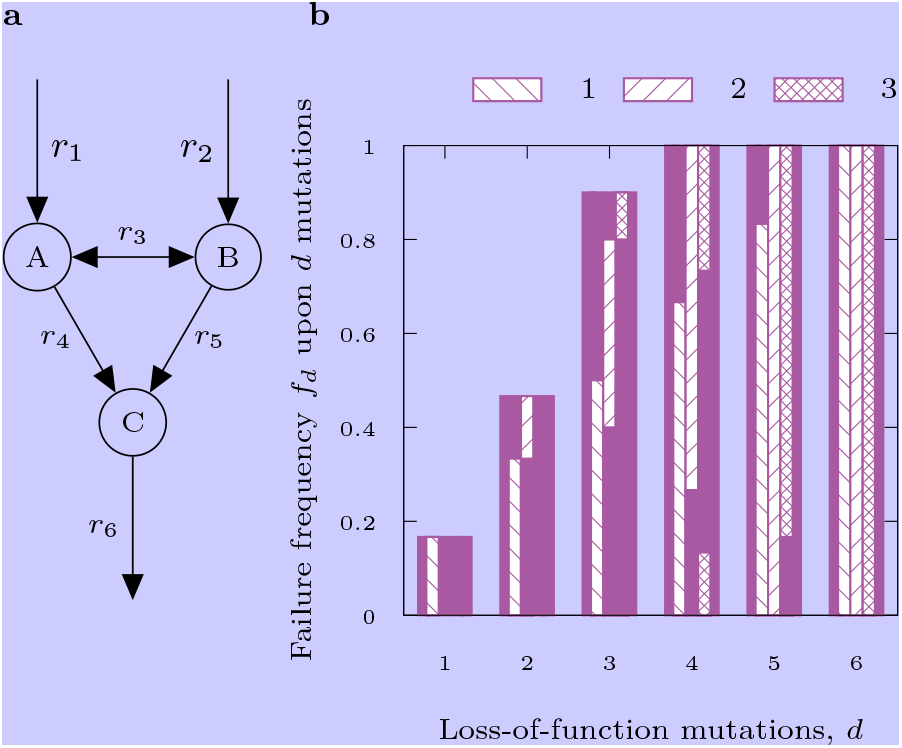
Toy network (a) and failure frequency as function of the number of LOF mutations (b). Hatched bars indicate the contributions of the essential reaction (*r*_6_, column 1), the two synthetic lethal pairs ({*r*_1_, *r*_2_} and {*r*_4_, *r*_5_}, column 2), and the two synthetic lethal triplets ({*r*_1_, *r*_3_, *r*_5_} an {*r*_2_, *r*_3_, *r*_4_}, column 3). Note the increasing overlap between the bars for increasing *d*, indicating that large parts of *f*_*d*_ can be estimated without knowledge of higher order combinations of mutations. For a complete list of all lethal combinations see **Supplementary Table 1**.

Here, we present fundamental improvements – both in theory as well as software implementation – of our PoF-calculating algorithm [13]. Provided with a list of low-cardinality MCSs and running on a personal computer, the program is now capable of determining the PoF of a GSMM at reasonable accuracy in under a minute, whereas hours of high-performance computing time have been required in the past. With the improved tool at hand we computed the PoF across a large panel of Enterobacteriaceae and 30 fungal species and show that their structural, metabolic robustness does not correlate with their ability to thrive in nutritionally diverse environments.

## TERMINOLOGY AND DEFINITIONS

The PoF, *F*, of a metabolic network with *r* reactions is defined as the probability that a given number of random loss-of-function (LOF) mutations disables growth (i.e. is lethal). It can be expressed as the probability-weighted average of the failure frequency

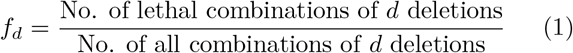

for LOF mutations in exactly *d* enzymes (reactions) being lethal. This gives

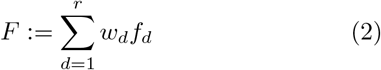

with the discrete binomial distribution

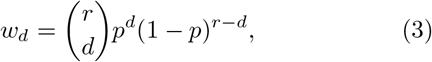

which — for a constant reaction-level mutation rate *p* — describes the probability that *d* LOF mutations occur.

Eq. (1) can be expressed in a computationally more approachable way using combinations of MCSs [13]. In this context, an MCS is a minimal set of reaction deletions that suppresses growth [12]. Essential reactions, synthetic lethal pairs or synthetic lethal triplets are examples of MCSs with cardinalities one, two, or three, respectively. In the Supplementary Notes we expand on [13] and show that, if all MCSs of a metabolic network are known, eq. (2) can be formulated as

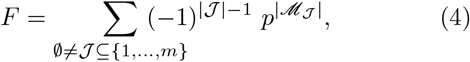

where ℳ_*𝒥*_ denotes the union ∪_*j*∈*𝒥*_ ℳ_*j*_ of one or more MCSs with the (multi)-index *𝒥* running over the power set of the indices {1, …, *m*} of all *m* MCSs. In other words, this means that every possible combination of one or more MCSs is a summand in eq. (4). In **Supplementary Table 2** we provide an evaluation of Eq. (4) for the toy network in **Fig. 1a**.

Note that according to eq. (4), *F* solely depends on the total number of MCSs, *m*, and the cardinalities of the members of their power set (i.e. on how the MCSs overlap with each other), but not on the number of reactions in the network. Thus, the PoF is independent of network size.

### Estimating *F* in large models

Eq. (4) determines the PoF exactly as long as (i) all *m* MCSs are known and (ii) the sum is evaluated over all their possible combinations (i.e. the full power set with 2^*m*^ *–* 1 members). For GSMMs both requirements cannot be met as (i) the enumeration of all high-cardinality MCSs in a large network is computationally intractable and (ii) even if it was not, the number of possible combinations would soon exceed the number of atoms in the universe. Thus, approximations are indispensable.

It is possible to systematically construct the first *m*′ MCSs of lowest cardinality [15], which have the greatest impact on the PoF. Hence, *F* can still be reliably estimated, even when only a relatively small number of MCS are known [13]. This addresses the first issue. Yet, even if the number of included MCSs is reduced from *m* to *m*′, the exponential explosion of the summands remains critical. In the simplest, least accurate case, we consider only essential reactions, i.e. all *m*_1_ MCSs with |ℳ_*j*_ | = 1, and eq. (4) simplifies to

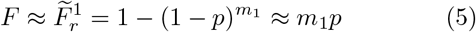

(for derivation see Supplementary Notes). When all MCSs up to a certain cardinality are known, we call this cardinality *d*_*m*_ (in this case *d*_*m*_ = 1). The number of essential reactions, *m*_1_, times the mutation rate, *p*, is in fact the leading term in the sum of eq. (4). To improve accuracy, we resort to eq. (2) and, rather than summing over all *r*, we truncate the sum to *d → d*_0_ so that

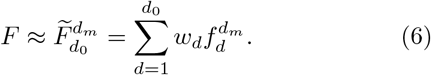

Here, the super-script *d*_*m*_ indicates that 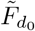 and *f*_*d*_ are computed with all MCSs up to cardinality *d*_*m*_. However, we show in the Supplementary Notes that the resulting error can be further reduced.

**Fig. 2** serves to illustrate the maximum error *ε*_max_ for 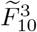 in the *E. coli* GSMM *i*JO1366 (*d*_*m*_ = 3; all MCSs with cardinalities smaller than 4 are known). For *d* ≤ *d*_*m*_ = 3 all 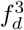 are exact. MCSs with cardinalities greater than three have not been calculated and their contributions are unknown. Therefore, in the worst case all cut sets of cardinality 4 could be lethal leading to 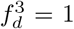 for *d* ≥ 4. The resulting maximum error is thus represented by the hatched bars in the inset of **Fig. 2**. It can be calculated via

**Fig. 2.**
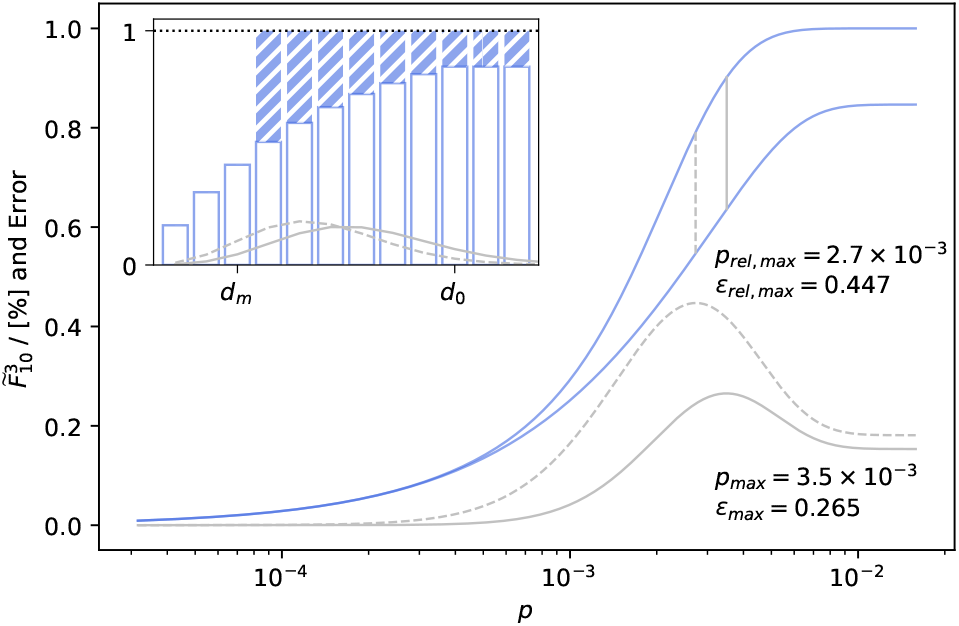
Maximum error of 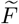. Lower 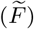 and upper 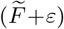 bounds (blue lines) as well as relative and absolute errors (dashed and solid grey lines) for the PoF of a GSMM of *E. coli* with *d*_*m*_ = 3 and *d*_0_ = 10 vs. LOF mutation rate *p*. Vertical grey lines indicate the position of the respective maximum value. The inset shows *f*_*d*_ for 1 ≤ *d* ≤ 12 (white bars) and the corresponding maximum error (blue bars) as well as the shape of the binomial distribution at the mutation rates with the greatest absolute and relative errors.

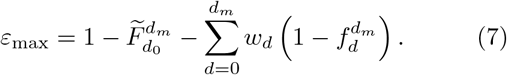

For *i*JO1366 in **Fig. 2** the maximum absolute and relative errors peak around *p* = 3 × 10^−3^ and quickly approach zero for *p <* 0.5 × 10^−3^. Note that the sum in eq. (7) is independent of *d*_0_.

## IMPLEMENTATION

In our previous work [13], we approximated the (total) PoF by computing *f* (*d*; *m*_0_) only for *d* ≤ *d*_0_ = *d*_*m*_ and formulated a parallelised algorithm to carry out the truncated sum. Here, we present `pof2.0` that includes further algorithmic improvements and allows us to quickly evaluate 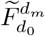 for larger *d*_0_, which has two implications: (i) For a given number of MCSs *F* can be predicted more precisely and (ii) to achieve certain accuracy the number of required MCSs can be reduced.

At the core of our method lies the fact that any lethal cut set must be a superset of at least one MCS. In order to estimate 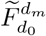, |ℳ _*𝒥*_| (the number of knocked out reactions) as well as | *𝒥*| (the number of MCSs in the cut set) need to be found for every possible combination that can be formed from the network’s known MCSs (i.e. their power set). Our program achieves this by a recursive procedure, the basis of which has been outlined in [13]. In short, we first sort the list of MCSs by cardinality. Then, for every element in the sorted list we iterate over all MCSs that appear earlier in the list and form the respective unions with |*𝒥*| = 2. If a union with |ℳ_*𝒥*_| ≤ *d*_0_ is not a subset of a previously encountered cut set, we again iterate over all elements in the list up to the earliest MCS already present in that union to form all cut sets with |*𝒥*| = 3 and so on. |ℳ_*𝒥*_| and |*𝒥*| of every lethal combination are recorded to be evaluated once recursion has finished. The actual implementation employs a few shortcuts that considerably reduce the number of recursions required.

Moreover, in metabolic pathway analysis it is common practice to compress networks before analysis [16]. This reduces the number of metabolites and reactions without losing information and is necessary for efficient utilisation of computational resources. Naturally, compressed networks also contain drastically fewer MCSs. `pof2.0` is capable of handling linearly compressed networks, which entails a major leap in performance. The code is available at www.github.com/julibeg/PoF.

## RESULTS

### PoF computes within seconds

To assess the computational efficiency of `pof2.0`, we analysed its performance as a function of the number of processed MCSs for *E. coli*’s central carbon metabolism model (CCMM) [17] and its GSMM *i*JO1366 [18] (**Supplementary Figure 2**). In both cases, we observed a considerable reduction in run-time cost per additionally processed MCS compared to the previous implementation [13]. For instance, after processing ∼2,000 MCSs of the CCMM, we reached a ∼100-fold speedup, the majority of which can be attributed to network compression. Extrapolating run-times to ∼796,070 MCSs (*d*_*m*_ = 15) gives an estimate of ∼280 hours for the previous implementation while `pof2.0` took less than a second. Due to these advances, the bulk of the computational burden in estimating structural robustness now clearly lies at MCS enumeration.

The preceding sections established the theoretical foundation and the computational feasibility of our approach to quantify structural robustness. Next we capitalise on the reduced run-time of `pof2.0` by analysing hundreds of GSMMs in two case studies in order to showcase the utility of our approach and examine the nature of metabolic robustness. Unless stated otherwise, all PoF approximations reported below were calculated with *d*_*m*_ = 3, *d*_0_ = 10, and *p* = 10^−4^.

### Case study I – robustness correlates with carbon source molar mass

To explore the influence of varying carbon sources on the robustness of growth in *E. coli* we computed the PoF of *i*JO1366 [18] growing on minimal medium under aerobic conditions for all its 181 single, growth-supporting carbon sources. As shown in **Fig. 3**, the PoF decreased with increasing molar mass of the carbon source (Pearson: *r* = –0.30, *p* = 3 × 10^−5^; Spearman: *r* = –0.48, *p* = 8 × 10^−12^).

**Fig. 3.**
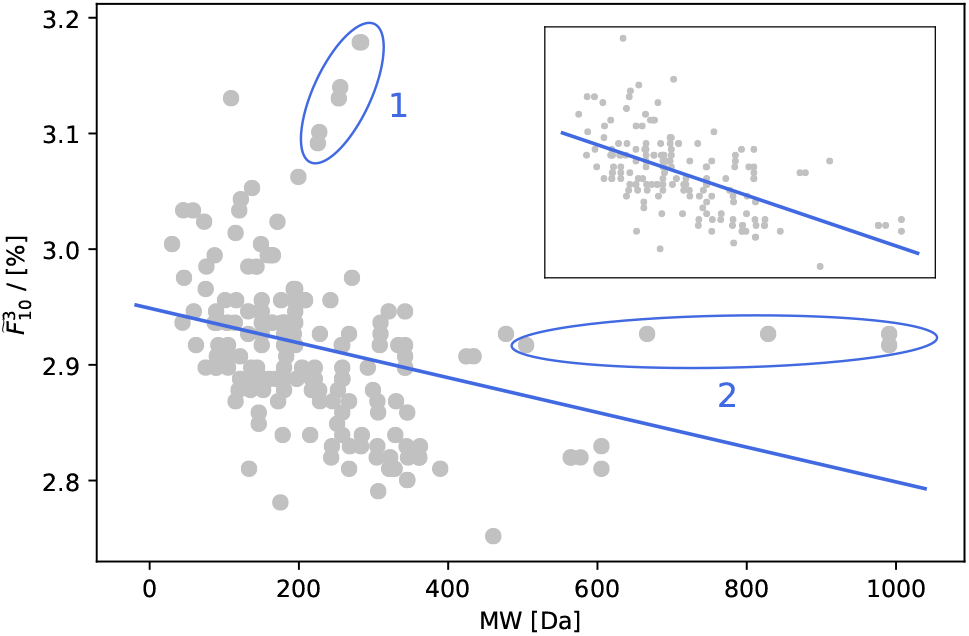
PoF vs. carbon source molar mass. 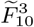 as function of the carbon source’s molar mass for all single, growth-supporting carbon sources in *E. coli i*JO1366. The line represents a linear fit to the data. Ellipses single out two spikes (see main text for details). The inset shows data with spikes removed and improved fit.

**Fig. 3** features two pronounced spikes (marked “1” and “2”). In spike 1 the PoF increases with the molar mass of the respective carbon sources, which are all medium- and long-chain fatty acids. While in reality the same enzymes process fatty acids of different lengths during beta-oxidation, these steps are represented as distinct reactions in metabolic models. Thus, contrary to *in vivo*, fatty acids are catabolised by increasingly longer linear reaction chains *in silico*. Spike 2, on the other hand, is comprised of cases where a dramatic increase in molar mass has hardly any effect on the PoF. Responsible are oligomaltoses (i.e. starch oligomers), most of which reach central metabolism via the same number of reactions *in silico*. Without the spikes, correlation increases to: Pearson - *r* = –0.57, *p* = 5.4 × 10^−16^; Spearman - *r* = –0.58, *p* = 1.3 × 10^−16^.

We further investigated the impact of the carbon source’s molecular composition on the PoF, which revealed a hierarchy based on chemical composition. On average, “pure” carbon sources (i.e. consisting only of C, H, and – possibly – O atoms) supported less robust growth than substrates containing either nitrogen or phosphate or both, see **Supplementary Figure 3**.

Growth on “elementally richer” substrates being more robust can be explained by higher redundancy. For instance, when growing on an amino acid, otherwise lethal mutations in ammonium transport and assimilation pathways would become irrelevant. As an example consider the 298 MCSs of cardinality 1 in the joint set of MCSs for aerobic growth on either glucose or glutamate, out of which 285 were identical for both. Only eight and five were specific to glucose and glutamate, respectively, which is reflected in a lower PoF for the latter. Three out of the eight essential reactions unique to growth on glucose are involved in ammonium transport. This illustrates how *F* is linked to topological differences in metabolic networks.

### Case study II – the origin of biological robustness

The evolutionary origin of genetic robustness – despite its ubiquity [2, 19] – remains strongly debated [20, 21]. Approaches to explaining its emergence can be roughly split into three categories: congruent, adaptive and intrinsic [20]. Contrary to the adaptive hypothesis, which states that genetic robustness is selected for directly [22–28], the congruent hypothesis postulates that it rather arises as a byproduct of selection for environmental resilience [29–31]. The intrinsic hypothesis, on the other hand, negates any evolutionary impact and argues that robustness is a fundamental property of complex systems and networks [32–34]. However, due to its multi-factorial nature, it is more likely that no single mechanism is responsible for the development of biological robustness. The actual origin may rather comprise a combination of the hypotheses introduced above.

To test the impact of the congruence mechanism (“environmental robustness breeds genetic robustness”) we speculated that organisms capable of growing in many different conditions (robust towards changes in the chemical environment) would exhibit low PoFs (robust towards LOF mutations). Accordingly, structural robustness in metabolic networks should correlate with the number of substrates enabling growth. To evaluate this idea, we calculated the PoF and counted the aerobic, growth-supporting carbon sources for GSMMs of 53 *E. coli* and *Shigella* [35], 406 *Salmonella* [36] as well as 30 fungal [37] strains.

### Robustness correlates with the diversity of growth media

For the *E. coli* and *Shigella* models the PoF was calculated twice – once with regard to growth on glucose minimal medium and once for minimal medium with all growth-supporting carbon sources enabled. Being allowed to grow on all carbon sources, the networks showed considerably higher robustness compared to growth on glucose alone (**Supplementary Figure 4**). In fact, when testing *i*JO1366, the average of *F* decreased monotonically with the number of carbon sources available (see **Supplementary Figure 5**). This confirms the naïve expectation that fatal reaction deletions are more strongly buffered in richer media. However, the magnitude of the two PoFs for a given model were correlated (Pearson: *r* = 0.77 *p* = 1 × 10^−11^; Spearman: *r* = 0.60, *p* = 3 × 10^−6^).

Similarly, we found growth on minimal glucose medium to be considerably more robust than optimal growth on the same medium (**Supplementary Figure 4a**). In fact, the distribution of the PoF across the models was quite consistent for the different conditions with the exception of *Shigella boydii* Sb227. This strain showed the greatest robustness among all *E. coli* and *Shigella* models at glucose-optimal growth while ranking last when all carbon sources were enabled (**Supplementary Figure 4b**).

### PoF does not correlate with carbon utilisation capabilities

**Fig. 4** shows the PoF across all *E. coli, Shigella, Salmonella* and fungal models versus the number of carbon sources that allow the respective networks to grow. Among the *E. coli* and *Shigella* strains it ranged from 2.78% (*E. coli* IAI39) to 2.95% (*S. boydii* CDC 3083-94) with an average of 2.87%. For *Salmonella* and the fungi it was considerably lower with an average of 2.62% and 1.04%, respectively. Within the respective groups there were no correlation between *F* and the number of carbon sources. On the contrary, most *E. coli* models sit on a straight horizontal line from 167 to 186 different carbon sources with the PoF being practically identical at 2.8685%. Similarly, out of 406 *Salmonella* models 213 have essentially the same 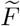 of 2.6059% and grow on 143 to 168 carbon sources. Finally, the fungi showed the highest amount of variability yet again no correlation with the number of carbon sources.

**Fig. 4.**
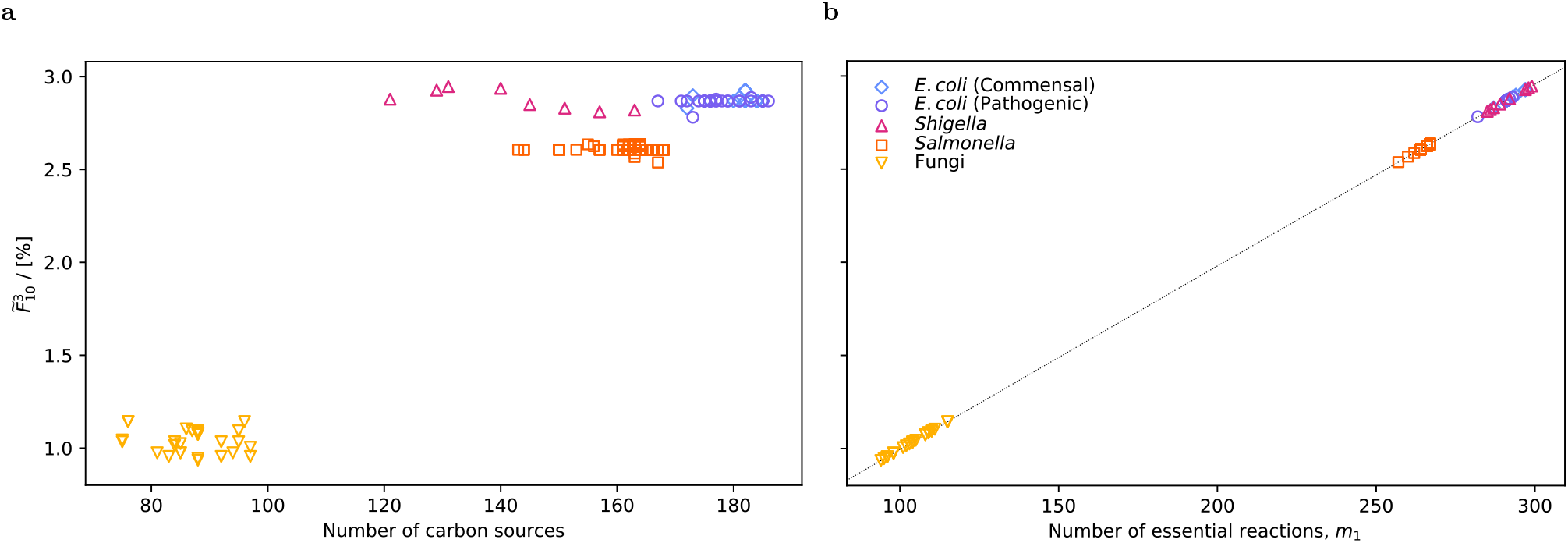
Genetic robustness vs. environmental robustness. **a** Estimated total PoF, 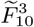, as a function of the number of growth-supporting carbon sources for all models tested. Growth on glucose minimal medium under aerobic conditions was simulated in GSMMs of various commensal and pathogenic *E. coli, Shigella*, and *Salmonella* strains as well as 30 fungal species. **b** 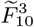 vs. *m*_1_ (i.e. number of essential reactions). The dotted line represents the PoF approximation using eq. (5) and the number of essential reactions alone.

The observation that many strains have (almost) identical PoF values, although differing substantially in carbon utilisation capability, suggest that additional carbon sources mostly only add extra uptake reactions to the network rather than increasing connectivity. For example, all *E. coli* models with 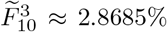 feature the same set of essential reactions, albeit some can grow on close to twenty more carbon sources compared to others. The linear fit in **Fig. 5** corroborates this assumption as the number of reactions increases by ∼3.28 for every additional carbon source. These three reactions usually consist of an exchange reaction for importing the carbon source across the model boundary and two transport reactions across the outer and inner membrane. Therefore, roughly only every third carbon source adds one non-transport reaction to the model leaving little room for increasing metabolic connectivity.

**Fig. 5.**
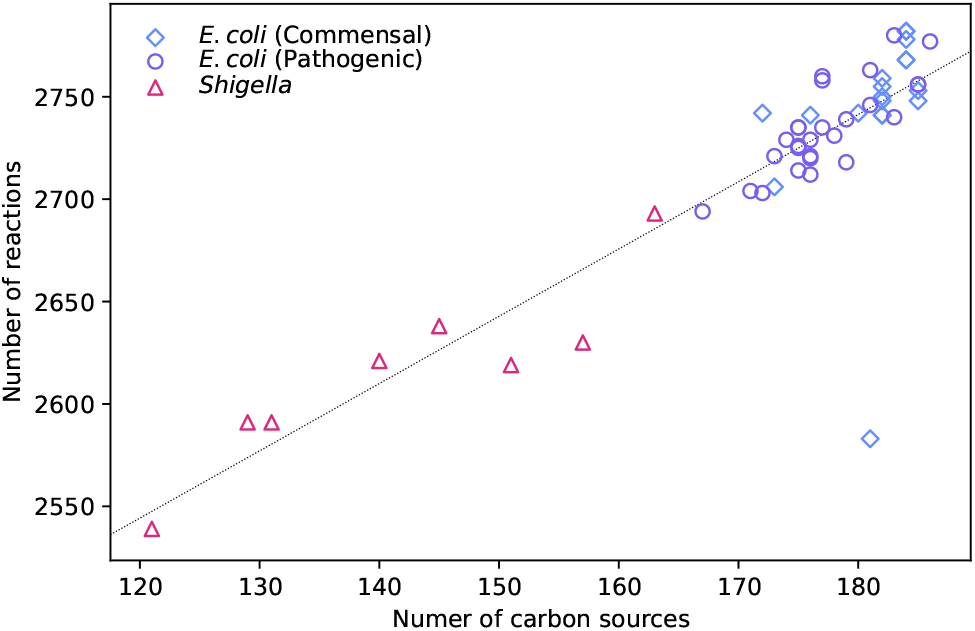
Reactions vs. C-sources. Number of reactions vs. number of carbon sources for all native *E. coli* and *Shigella* models. The slope of the linear fit (dashed line) is ∼3.28.

To further examine this hypothesis we computed the size of *E. coli*’s “shared reactome”, i.e. the set of reactions present in all *E. coli* GSMMs. Under the investigated conditions (aerobic growth on glucose minimal medium) it was 1, 336*/*1, 774 = 75%. Considering only the *E. coli* strains with virtually equal PoF it increased to 1, 447*/*1, 772 = 82%, indicating that the PoF is a good predictor for network overlap. Similarly, a pairwise comparison across all *E. coli* and *Shigella* models revealed that the difference in PoF correlates with the pairwise network dissimilarity, 1 – *σ*_*i,j*_, of any two networks *M*_*i*_ and *M*_*j*_ (see **Fig. 6**). Here, dissimilarity was defined as the complement of the pairwise network similarity

**Fig. 6.**
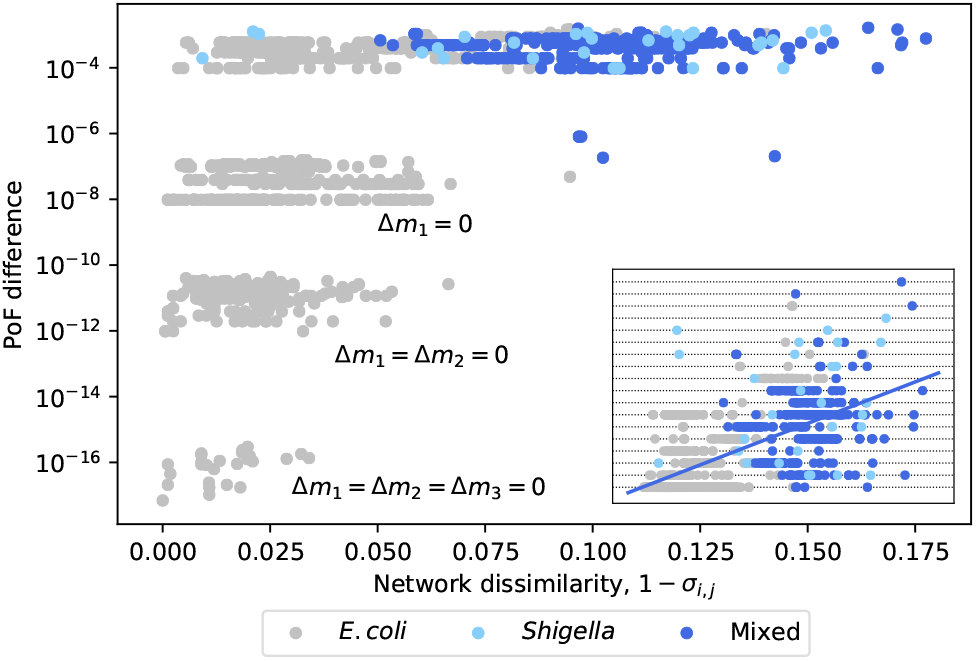
Pairwise difference in the PoF across all *E. coli* and *Shigella* GSMMs vs. network dissimilarity. Strains with different *m*_1_ cluster around 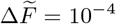; strains with equal *m*_1_ around 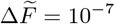; equal *m*_1_ and *m*_2_ around 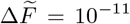; equal *m*_1_, *m*_2_, and *m*_3_ around 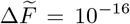. Mixed pairs of strains are in blue; pairs of *E. coli* and *Shigella* strains are in grey and light blue, respectively. Note the logarithmic scale on the y-axis. The inset shows the same data in linear scale and a linear fit. Horizontal dotted lines correspond to Δ*m*_1_.

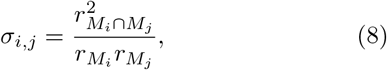

Where 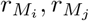, and 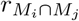 denote the number of reactions in the models *M*_*i*_, *M*_*j*_, and of those found in both, respectively.

In addition to greater robustness the *Salmonella* models also displayed reduced nutritional flexibility; being able to grow on only 164 carbon sources on average as opposed to *E. coli*’s 179. Moreover, their relative shared reactome was slightly larger at 1274*/*1628 = 78%.

With only 1383*/*2153 = 64% of reactions belonging to the shared reactome, the fungal models showed larger diversity. This is not surprising, since the group of fungal models featured 30 species from different genera as opposed to multiple strains / species from the same genus for the bacterial models.

In spite of decreased nutritional flexibility all fungal species showed considerably larger robustness as opposed to the bacterial models discussed above. However, it appears that this observation is an artefact of the different model reconstruction procedures rather than a genuine biological effect. The fungal models’ biomass consisted of 44 components, while the bacterial biomass reactions were considerably more detailed (51 and 68 components for *Salmonella* and *E. coli*/ *Shigella*, respectively) including additional vitamins and metal ions. Consequently, the growth medium was more complex for the bacterial models (containing 18 components) as opposed to the fungal models with only five components (see **Supplementary Table 4**). Moreover, the more complex the biomass composition, the more likely it is effected by LOF mutations. In fact, when using the fungal biomass reaction as objective in *i*JO1366, the PoF decreased from 2.89% to 1.53% (**Supplementary Figure 6**). Thus, as the biomass composition is different for models of the respective reconstruction groups (fungal, *E. coli*/*Shigella*, and *Salmonella*) we conclude that comparisons within groups are appropriate, but not across groups.

When comparing pathogenic with non-pathogenic models within the same reconstruction group, no differences in the PoF have been found (**Supplementary Figure 7**). However, given the small number of purely pathogenic strains/species and the lacking genetic diversity among them, general trends cannot be inferred.

### Essential reactions dominate metabolic robustness

For the models assessed here, a comparison with eq. (5) (right panel in **Fig. 4**) reveals that even at the relatively high *p* = 10^−4^, the number of MCSs with cardinalities two and three was not large enough to contribute noticeably to 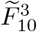. This observation – under the assumption that it can be generalised to most bacterial models growing on minimal medium – suggests that a reasonably accurate approximation for the structural robustness of these networks could be obtained solely from the number of essential reactions, *m*_1_, which is easy to compute.

**Fig. 6** further highlights the minuscule contribution of the MCSs with |ℳ_*j*_ | ∈ {2, 3} as all pairwise PoF-differences between two models with the same number of essential reactions are smaller than 10^−6^. Similarly, the PoF differences among pairs of models with the same number of essential reactions as well as synthetic lethal pairs (*m*_1,*i*_ = *m*_1,*j*_ and *m*_2,*i*_ = *m*_2,*j*_) lie between 10^−12^ and 10^−10^ and are mostly determined by differences in the number of synthetic lethal triplets Δ*m*_3_ (**Supplementary Figure 8**).

As *F* is almost exclusively determined by *m*_1_ we identified the essential reactions shared by all *E. coli* and *Shigella* models and the corresponding KEGG [38] pathways (**Fig. 7**). Interestingly, (central) carbon metabolism only hosts a small number of essential reactions and thus constitutes a minor contribution to the PoF. Instead, amino acid synthesis appears to be buffered by redundancy to a lesser extent.

**Fig. 7.**
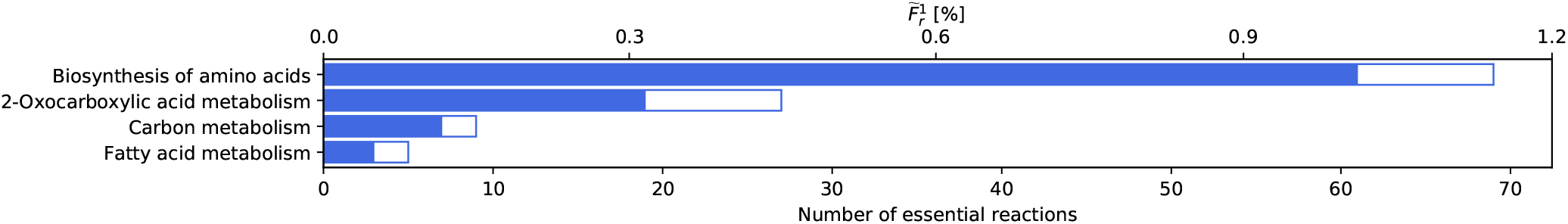
Essential reactions grouped by pathway. Number of essential reactions grouped by KEGG pathways occurring in (empty bars) and shared by (solid bars) all *E. coli* and *Shigella* models.

## DISCUSSION

Robustness is a key feature of biological systems. Yet, its exact quantification remains challenging. In a previous work, we have modeled the robustness of metabolic networks by counting the frequency of the binary network response (as measured by growth or no growth under steady-state conditions) upon random reaction deletions [13]. Similar mathematical strategies are commonly used in engineering to evaluate the reliability of safety-critical structures, for example in aircraft [39]. Here, we expanded on this approach and presented a comprehensive theoretical description which we validated by confirming the buffering capabilities of diverse growth media [40, 41].

Capitalising on reduced run-times for GSMMs we were able to evaluate the PoF across 459 members of the family Enterobacteriaceae (amongst them commensal and pathogenic *E. coli, Shigella* as well as *Salmonella* strains) and 30 species of fungi growing on glucose minimal medium under aerobic conditions. Although the closely related *E. coli* and *Salmonella* strains, whose lineages separated 140 million years ago [42], share a large fraction of their genetic material [43, 44], we found the *Salmonella* models to be considerably more robust. In fact, *Salmonella* serovars were shown to have fewer essential genes (relative to the genome size) than *E. coli* MG1655 [45–47], which is also reflected in the corresponding metabolic models. This indicates higher redundancy and supports our *in silico* analysis of the respective GSMMs. However, in this work we have not undertaken any phylogenetic analyses and thus cannot discuss evolutionary trajectories that could explain these differences.

Compared to the bacterial models the fungal species analysed here showed drastically higher robustness. However, it should be stressed that, if GSMMs that have been generated via different reconstruction pipelines are compared, caution has to be exercised as differences in the reconstructions process add to the observed PoF. For instance, the biomass composition is more detailed in the Enterobacteriaceae models as they require metal ions in their minimal *in silico* medium to support growth, whereas the fungal models do not. This prohibits a direct comparison between the respective groups. In general, we observed an increase in the PoF with the complexity of the biomass formulation. This is consistent with earlier reports that the size of essential networks scales with the number of biomass components [48]. However, to turn this drawback into a benefit, the PoF could be used as a quality indicator. For instance, we were able to track the smaller PoF of *E. coli* IAI39 compared to the average PoF of the *E. coli* strains back to a lumped reconstruction of folate biosynthesis.

Within the groups of models that share the same reconstruction pipeline (*E. coli*/ *Shigella, Salmonella*, and fungi), our findings do not support a connection between nutritional flexibility and cellular robustness. In line with this result, the diversity of genetic capabilities observed in *Salmonella* was shown to not be indicative of strain-specific lifestyle [36]. Here we find that this is true not only for Enterobacteriaceae but also for fungi. Although the strain-specific portions of the pan-reactome are largely associated with altered carbon metabolism [35, 36], our analysis reveals that this does not lead to increased connectivity (**Fig. 5**) – and thereby structural robustness – of the respective network. This reflects the notion that metabolism is organised in a bow tie structure with a large number of diverse nutrients being ‘fanned into’ a tightly knotted core network from where just a few key metabolites ‘fan-out’ again to produce all building blocks and macromolecular compounds that make up a cell [49, 50]. The observation, that on average the network size of *E. coli* and *Shigella* GSMMs increased by only three reactions for every additional carbon source (**Fig. 5**) – a number that is in good agreement with the previously derived two reactions per carbon source for minimal metabolic models [48] – also corroborates this interpretation.

The congruent hypothesis has recently also been challenged by Ho and Zhang [21]. Their analysis was based on the reallocation of growth-optimal flux distributions in a single GSMM of *E. coli* MG1655 (computed via minimisation of metabolic adjustments [51]) upon genetic and environmental perturbations [21]. Due to the use of an optimisation principle, their analysis was inherently biased, while our approach is more general in the sense that it takes the complete network structure (i.e. all possible flux re-routings) into account [52]. It could be argued that robustness is most relevant in the fittest states. However, our results show that structural robustness impacts all levels of growth-optimality in a similar way (**Supplementary Figure 4**).

Classically, an unbiased characterisation of metabolic networks is achieved in terms of minimal functional units that are able to characterise all feasible steady-state flux distributions [6, 53–55]. Alternatively, a network can be equally well described by failure modes, called MCSs [11]. This is mirrored by the fact that, according to eq. (4), the PoF is solely determined by the MCSs and independent of the network’s size. Thus, it is a true feature of a network’s structural topology.

In principle, *F* can be computed exactly. However, in practice it suffers from (i) the combinatorial explosion of the number of MCSs in GSMMs and (ii) the exponential explosion of the number of summands in eq. (4) (i.e. the power set of the MCSs). Nonetheless, evaluating the PoF remains feasible as the (relatively few) low-cardinality MCSs are most dominant [13]. In fact, for many models growing on minimal media the number of essential reactions already suffices. This is due to the fact that it is far more likely that a randomly selected set of *d* deletions contains e.g. one or more essential reaction(s) than a complete MCSs of cardinality *d*. Hence, high-cardinality MCSs are not as relevant and can be neglected provided *d*_0_ is sufficiently large and *p* is sufficiently small (**Fig. 2**).

The probability of LOF mutations in a given number of reactions follows the binomial distribution, consequently depending on the total number of reactions in the model as well as the per-reaction mutation rate *p*. As we reported solely comparative analyses here, we focused on relative differences in *F* and chose *p* to be 10^−4^ per gene. In order to correctly calculate absolute values, a more accurate, strain-specific estimate for *p* would be required. However, quantifying deleterious mutation rates proves challenging, as LOF mutations with strong effects are usually cleared from populations quickly, thereby evading experimental detection. Kibota and Lynch [56] give a lower bound of 2 × 10^−4^ deleterious mutations per genome and generation of *E. coli*, while estimates for the general mutation rate are in the range of 5 × 10^−3^ to 10^−4^ per genome and generation [57–60]. Hence, the rate for LOF mutations of a given gene (or reaction) can be confidently assumed to be very low.

This has implications for the computation of *F*, as fewer MCSs are required to give precise estimates in case of smaller mutation rates. For instance, in a network with *r* = 3000 and 300 essential reactions, the maximum relative error of the analytical PoF estimate from the essential reactions alone would be around 1% with *p* = 10^−6^ (see **Supplementary Figure 1c**). Thus, in such a case performing recursion would not be necessary.

In fact, for all models analysed here the number of essential reactions dominated 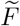, enabling accurate predictions with eq. (5) even at the unrealistically high *p* = 10^−4^. Thus, genetic robustness could simply be estimated given the number of essential reactions or genes.

Due to the binary nature of LOF mutations underlying the concept of the PoF, it can only measure structural redundancy in metabolic networks by analysing their topology in the steady state. Alternative forms of resilience or robustness, for instance absolute concentration robustness [61] or regulatory feedback mechanisms, are not addressed. Moreover, due to its focus on the reaction level, the PoF disregards some genetic information (e.g. about the presence of duplicated genes coding for the same enzyme). However, we have demonstrated previously that PoF analyses at the level of genes and at the level of reactions yield qualitatively similar results [13]. Incidentally, recent advances in software for the enumeration of genetic MCSs [62, 63] allow for extending the scope of our method to the gene level. Additionally, we have shown here that the PoF is independent of the network’s size and mostly determined by the number of essential reactions (or genes, for that matter). Thus, given knowledge of the essential genes [64] (and potentially the synthetic lethal pairs, which have been determined for some model organisms like yeast [65]), eqs. (4) and (5) enable us to evaluate the PoF on the genome level without the need for a metabolic reconstruction altogether. When relying only on the essential genes for PoF estimation, the theory simplifies greatly. Therefore, in future work the theoretical framework presented here could be expanded such that ways for incorporating gene lengths in the analysis could easily be devised, which would further improve the accuracy and reliability of our approach for quantifying genetic robustness.

## METHODS

### Model acquisition and preprocessing

54 GSMMs of *E. coli* and *Shigella* strains [35] were downloaded from the BIGG Models database, version 1.6 [66, 67]. Additionally, 408 *Salmonella* GSMMs [36] (accession ID: MODEL1807280001) and 56 GSMMs of fungal species [37] (accession IDs: MODEL16042800{00-55}) were retrieved from BioModels [68]. The minimal media can be found in **Supplementary Table 4**. Native model sizes were ∼2,600 reactions for the bacterial and ∼6,700 reactions for the fungal species. To remove any artefacts that might have emerged in the automated model-reconstruction process, all models were made consistent with respect to growth on glucose. This means that every reaction unable to carry any flux when growing on the minimal medium (as determined by flux variability analysis [69]) was removed from the respective model. This reduced model sizes to ∼1650 reactions for *E. coli*/ *Shigella*, ∼1550 reactions for *Salmonella*, and ∼1750 reactions for the fungi. Note that in terms of the PoF analysis, “consistent” and “native” models are equivalent as *F* is independent of the number of reactions and the set of MCSs is the same for both. However, the models were made consistent nonetheless as it is commonly considered good practice and also reduces the computational load on MCS enumeration. Additionally, the *E. coli* and *Shigella* batch was also made consistent with regard to optimal growth on glucose, removing every reaction not participating in flux patterns providing the highest possible growth rate. Thereby, model sizes were reduced to ∼500 reactions. To avoid introducing any bias in the process of making the models consistent, glucose levels in the media were adjusted on a per-model basis beforehand so that all models would achieve the same growth rate. Moreover, for reasons stated below, the default upper and lower bounds on reaction fluxes were relaxed to *±*10, 000. The 54 *E. coli* and *Shigella* models contained 10 auxotrophs, one of which was capable of growing on its supplementary medium component as sole carbon source. It was therefore removed from the dataset. *Salmonella* featured 27 auxotrophs, two of which were able to grow on their supplements alone and were subsequently excluded from the analysis. The supplements required by the other auxotrophs were added to their respective media and can be found in **Supplementary Table 5** and **6**. Out of the 56 fungal models 26 were unable to grow in the specified conditions and thus dropped. The remaining 30 species (**Supplementary Table 3**) underwent additional pre-processing steps in order to harmonise reaction bounds prior to making the models consistent. All preprocessing was done in COBRApy v.0.17.0 [70].

### Computation of MCSs and PoF

To enumerate the models’ MCSs, the relevant information required was extracted from the respective SBML [71] or JSON files and reformatted accordingly. The data was then subjected to

i. Linear compression with an in-house Perl script. The target (i.e. biomass generation) reaction was exempt from compression.
ii. Transferring the MILP problem inferred from the compressed model into its dual form; again by an in-house Perl script.
iii. Solving the dual problem for MCSs with cardinalities up to 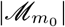 with an in-house implementation of the algorithm described in [15].

The compressed MCSs were then used for PoF evaluation. For all models, the loss-of-function mutation rate *p* was set to 10^−4^ per reaction.

### Single carbon source comparison for *E. coli*

Every carbon source available in *i*JO1366 was added separately to the minimal medium (**Supplementary Table 4**) bar glucose and the corresponding PoF with 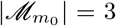 and *d*_0_ = 10 was determined. To reduce the computational burden in the MCS-enumeration step, the respective model was made consistent for every substrate. Again, in order to prevent introducing bias, the carbon source availability in the medium was tuned to allow for the same growth rate as glucose. This was infeasible for some carbon sources with very low yields due to internal model constraints (i.e. flux bounds). Therefore, these constraints were relaxed to *±*10, 000.

### Benchmarking PoF-implementations

For both *E. coli*’s CCMM [17] and the GSMM [18], low-cardinality MCSs were enumerated up to 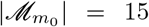 and 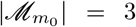, respectively. Then, for a range of *m* ≤ *m*_0_, the first *m* compressed MCSs were extracted and evaluated for all *d*_0_ ≤ 15 (CCMM) and *d*_0_ ≤ 8 (GSMM). Additionally, the MCSs were decompressed and evaluated again in the uncompressed form with both `pof2.0` and the old implementation available at https://github.com/mpgerstl/networkRobustnessToolbox. Computation was performed on an Intel^®^ Xeon^®^ CPU E5-2650 v3 @ 2.30GHz, and run times were recorded using Bash’s [72] built-in `time` command.

### Computation of the PoF

`pof2.0` was written in C++11 [73]. The implementation relies on Boost [74] for calculating binomial coefficients and uses Luigi Pertoldi’s progress bar [75]. The code is available at www.github.com/julibeg/PoF.

## AUTHOR CONTRIBUTION

Conceptualisation: J.L.E., J.Z. and M.P.G.

Methodology and Formal Analysis: J.L.E. and J.Z.

Data Acquisition and Curation: B.C. and J.L.E.

Software Implementation: J.L.E.

Visualisation: J.L.E. and J.Z.

Writing – Original Draft: J.L.E. and J.Z.

Writing – Review & Editing: all

Funding Acquisition: J.Z.

## DATA AVAILABILITY

All models used for analysis are publicly available [35–37]. *E. coli* and *Shigella* GSMMs can be obtained from the BIGG database [66, 67] (http://bigg.ucsd.edu) using the model IDs in Supplementary Data File 1. *Salmonella* GSMMs are available on BioModels [76] (https://www.ebi.ac.uk/biomodels) with the accession ID MODEL1807280001. Their individual IDs are listed in Supplementary Data File 2. Fungal models are available on BioModels as well. The respective accession IDs can be found in **Supplementary Table 3**.

## CODE AVAILABILITY

Source files for both, old and new, implementations of the PoF calculator are available at https://github.com/mpgerstl/networkRobustnessToolbox and www.github.com/julibeg/PoF, respectively.

## ACKNOWLEDGEMENTS

This work has been supported by acib. The COMET center: acib – Next Generation Bioproduction is funded by BMK, BMDW, SFG, Standortagentur Tirol, Government of Lower Austria and Vienna Business Agency in the framework of COMET - Competence Centres for Excellent Technologies. The COMET-Funding Program is managed by the Austrian Research Promotion Agency FFG.

