## Supplementary material for "Environmental flexibility does not explain metabolic robustness"

(Dated: October 4, 2020)

### CONTENTS

List of Figures 1

List of Tables 1

Supplementary Notes 2
- Derivation of eq. (4) 2
- Derivation of eq. (5) 3
- Improving the approximation of eq. (6) 4
- Derivation of eq. (7) 5

Supplementary Figures 6

Supplementary Tables 14

References 18

### LIST OF FIGURES

**1** Maximum error in the probability of failure (PoF) estimate. 6
**2** Speedup achieved by `pof2.0`. 7
**3** PoF vs. carbon source molecular weight grouped by elemental composition. 8
**4** PoF and nutritional restriction. 9
**5** PoF as function of the number of utilised carbon sources. 10
**6** Impact of biomass composition on the PoF. 11
**7** PoF of pathogens and non-pathogens. 12
**8** Pairwise PoF differences for all *E. coli* pairs with equal $\boldsymbol{m_1}$ and $\boldsymbol{m_2}$ 13

### LIST OF TABLES

**1** List of loss-of-function (LOF) mutations that result in cell death for the toy network in **Fig. 1a**. 14
**2** Evaluation of eq. (4) for the toy network in **Fig. 1a**. 15
**3** Fungal BioModels [1] accession IDs 16
**4** Minimal media. 16
**5** Auxotrophic *E. coli* and *Shigella* strains and the respective supplements required for growth. 17
**6** Auxotrophic *Salmonella* strains and the respective supplements required for growth. NMN, Nicotinamide mononucleotide 17

### SUPPLEMENTARY NOTES

#### Derivation of eq. (4)

In the following we expand on our previous work [2] and simplify eq. (2). Previously we showed that

$$f_d = \binom{r}{d}^{-1} \sum_{\emptyset \neq J \subseteq \{1,\ldots,m\}} (-1)^{|J|-1} \binom{r-|\mathscr{M}_\mathcal{J}|}{d-|\mathscr{M}_\mathcal{J}|}. \tag{1}$$

By inserting eqs. (1) and (3) into (2) and rearranging the sums, we obtain

$$F = \sum_{\emptyset \neq J \subseteq \{1,\ldots,m\}} (-1)^{|J|-1} \sum_{d=1}^{r} \binom{r-|\mathscr{M}_J|}{d-|\mathscr{M}_J|} p^d (1-p)^{r-d}. \tag{2}$$

For $d < |\mathscr{M}_J|$ the binomial coefficient is zero. Hence, the above equation becomes

$$F = \sum_{\emptyset \neq J \subseteq \{1,\ldots,m\}} (-1)^{|J|-1} \sum_{d=|\mathscr{M}_J|}^{r} \binom{r-|\mathscr{M}_J|}{d-|\mathscr{M}_J|} p^d (1-p)^{r-d}. \tag{3}$$

Finally, we introduce $i = d - |\mathscr{M}_J|$

$$F = \sum_{\emptyset \neq J \subseteq \{1,\ldots,m\}} (-1)^{|J|-1} p^{|\mathscr{M}_J|} \sum_{i=0}^{r-|\mathscr{M}_J|} \binom{r-|\mathscr{M}_J|}{i} p^i (1-p)^{r-|\mathscr{M}_J|-i} \tag{4}$$

and evaluate the sum over the full binomial distribution (which is equal to 1). Thus, we simply end up with eq. (4),

$$F = \sum_{\emptyset \neq J \subseteq \{1,\ldots,m\}} (-1)^{|J|-1} \, p^{|\mathscr{M}_J|}. \tag{5}$$

**Derivation of eq. (5)**

Suppose we know all $m_1$ essential reactions (minimal cut sets (MCSs) of cardinality 1) of a metabolic network. Then, eq. (4) becomes

$$\widetilde{F}^1 = \sum_{\emptyset \neq J \subseteq \{1,\ldots,m_1\}} (-1)^{|J|-1} \, p^{|\mathscr{M}_J|} \tag{6}$$

running over the power set of all $m_1$ MCSs. As those contain only one reaction each, the number of subsets with exactly $i$ MCSs is given by the number of combinations $\binom{m_1}{i}$. Moreover, as no MCS of cardinality 1 has a common element with any other MCS, we can compute the cardinality of any union of MCSs by counting the number of elements. Thus, the sum above simplifies into

$$\widetilde{F}^1 = -\sum_{i=1}^{m_1} \binom{m_1}{i} (-p)^i = 1 - \sum_{i=0}^{m_1} \binom{m_1}{i} (-p)^i = 1 - (1-p)^{m_1}, \tag{7}$$

where we first complete the binomial expansion and then used the binomial identity. Note that this expression also considers the impact of all possible combinations of reaction deletions containing at least one essential reaction. As higher-cardinality MCSs only increase $F$ further, eq. (7) represents a lower bound to the PoF.

**Improving the approximation of eq. (6)**

The PoF is given by

$$F = \sum_{d=1}^{r} w_d f_d \tag{8}$$

with

$$w_d = \binom{r}{d} p^d (1-p)^{r-d}. \tag{9}$$

Previously [2], we estimated $F$ by truncating the sum over $d$ from $r$ to $d_0$. However, $f_d$ increases monotonically with $d$, as any (higher-order) super-set that includes a MCS will also be lethal. Thus, any $f_d$ for $d \geq d_0$ will be at least as large as $f_{d_0}$ and eq. (8) can be approximated by (see **Supplementary Figure 1a**)

$$F \approx = \sum_{d=1}^{d_0} w_d f_d^{d_m} + f_{d_0}^{d_m} \sum_{d=d_0+1}^{r} w_d.$$

For very large $d$, $f_{d_0}^{d_m}$ might be smaller than the basal contributions of essential reactions, $f_{d_0}^1$. Thus by appropriately splitting the last sum and replacing $f_{d_0}$ by $f_d^1$

$$F \approx \widetilde{F}_{d_0}^{d_m} = \sum_{d=1}^{d_0} w_d f_d^{d_m} + f_{d_0}^{d_m} \sum_{d=d_0+1}^{\rho} w_d + \sum_{d=\rho+1}^{r} w_d f_d^1, \tag{10}$$

the approximation of the PoF can be further improved. Here, the second sum is carried out over all $d_0 + 1 \leq d \leq \rho$ where $f_{d_0}^{d_m} \geq f_d^1$ is a more accurate estimator for the true $f_d$ than $f_d^1$.

**Derivation of eq. (7)**

We assume that all MCS up to cardinality $d_m = 3$ have been computed, see **Supplementary Figure 1a**. Thus, all $f_d^{d_m}$ up do $d = 3$ are exact. At $d > 3$ we miss the contributions of higher-order MCSs. In the worst case all cut sets of cardinality four could already be lethal leading to $f_d^3 = 1$ for $d \geq 4$. The resulting maximum error is given by the weighted, cumulative sum of the hatched bars in **Supplementary Figure 1a** and can be computed by the compliment of the PoF estimate, $1 - \widetilde{F}_{d_0}^{d_m}$ minus the cumulated and weighted compliment of the error-free $f_d^{d_m}$s up to $d_m$,

$$\varepsilon_{\max} = 1 - \widetilde{F}_{d_0}^{d_m} - \sum_{d=0}^{d_m} w_d \left(1 - f_d^{d_m}\right). \tag{11}$$

In the worst case, i.e. if only essential reactions are known, eq. (11) simplifies into

$$\varepsilon_{\max}^1 = (1-p)^{m_1} - (1-p)^{r-1}\left[1 + p(r - m_1 - 1)\right]. \tag{12}$$

When added to eq. (7), the resulting expression gives an upper bound for $F$ (**Supplementary Figure 1**). We note that the ratio

$$\frac{\varepsilon_{\max}^1}{1 - \widetilde{F}^1} = 1 - (1-p)^{r-m_1-1}\left[1 + p(r - m_1 - 1)\right], \tag{13}$$

which represents the error relative to the network's robustness, only depends on the mutation rate $p$ and the number of non-essential reactions, $r - m_1$, see **Supplementary Figure 1c**.

### SUPPLEMENTARY FIGURES

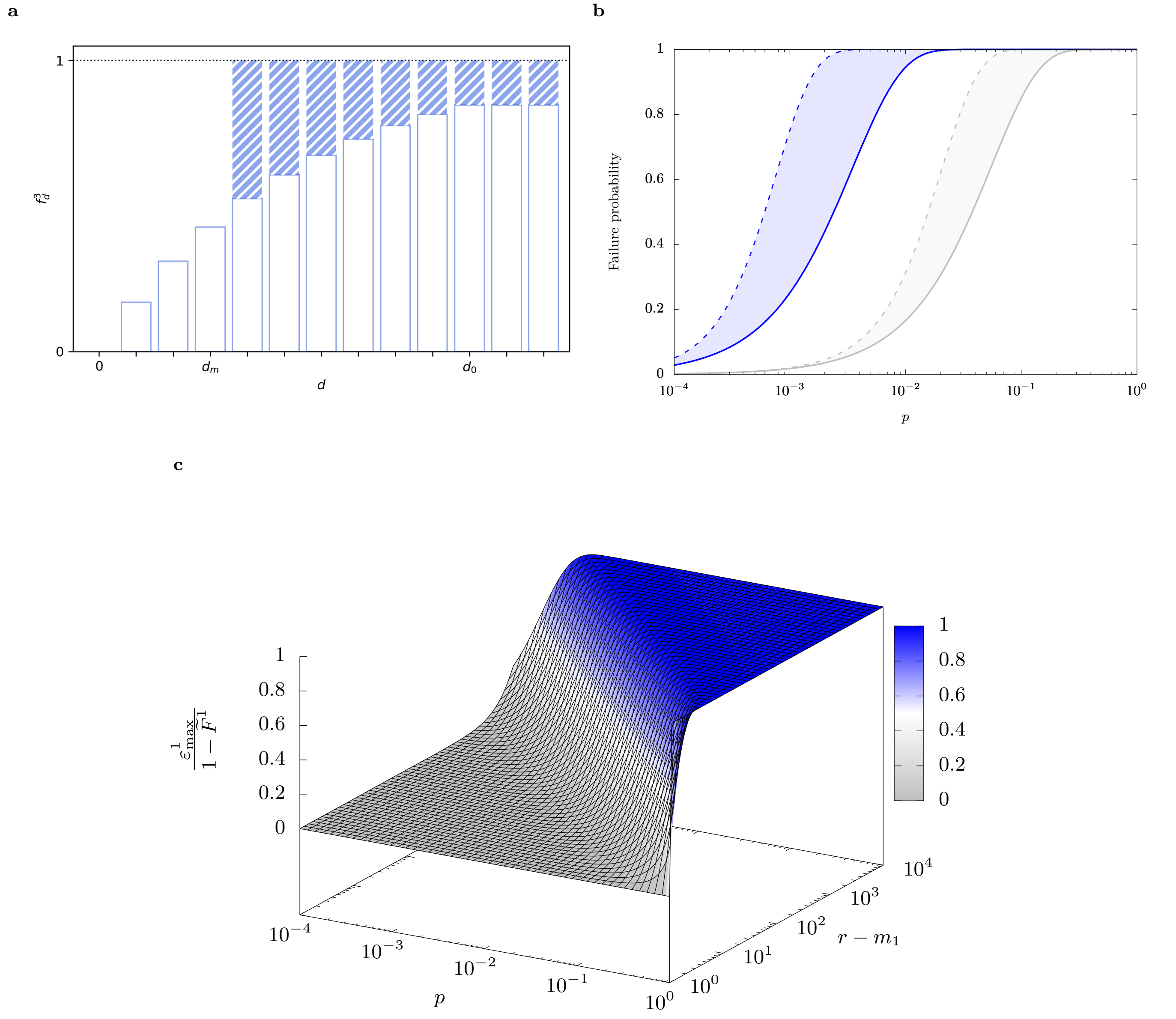


**Supplementary Figure 1**. **Maximum error in the PoF estimate. a** Visualisation of the maximum possible error (hatched bars) as function of $d$. Illustrated is a case with $d_m = 3$ and $d_0 = 10$. Note that for $d > d_m$, $f_d^3$ (empty bars) is only given by MCSs with cardinality up to 3. For $d > d_0$, we no longer update $f_d^3$, but use the value at $f_{d_0}^3$. **b** Lower ($\widetilde{F}^1$, solid lines) and upper ($\widetilde{F}^1 + \varepsilon^1_{\max}$, dashed lines) bounds of the PoF as function of the mutation rate $p$ given the number of essential reactions $m_1$ for two *E. coli* models. The GSMM *i*JO1366 [3] with $r = 2583$ and $m_1 = 289$ is depicted in blue; a model of *E. coli*'s CCMM [4] ($r = 95$, $m_1 = 18$) in grey. The shaded area indicates maximal possible uncertainty in $F$. **c** $\varepsilon^1_{\max}/(1-\widetilde{F}^1)$ according to eq. (13) as function for $p$ and $r - m_1$. The relative error grows with mutation rate and network size.

**a** **b**

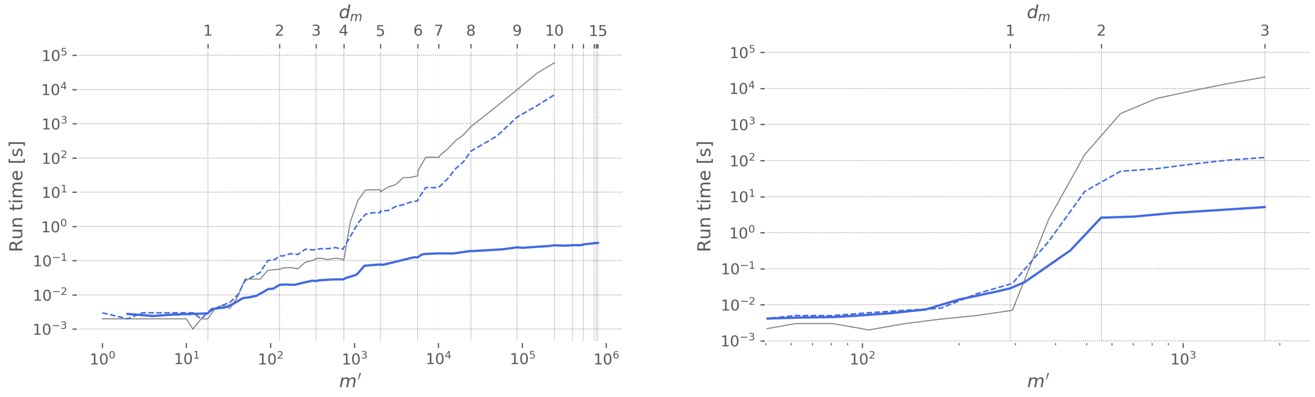


**Supplementary Figure 2**. **Speedup achieved by `pof2.0`.** Run time as function of the number of processed MCS, $m_0$, for `pof2.0` (blue) compared to the previous implementation (gray) [2]. We evaluated the total PoF for the CCMM of *E. coli* [4] with $d_m = 10$ and $d_0 = 15$ **a** as well as for the GSMM *i*JO1366 [3] with $d_m = 3$ and $d_0 = 8$ **b** simulating aerobic growth on glucose minimal medium. The blue solid and dashed lines illustrate performance of `pof2.0` for the linearly compressed and uncompressed networks, respectively.

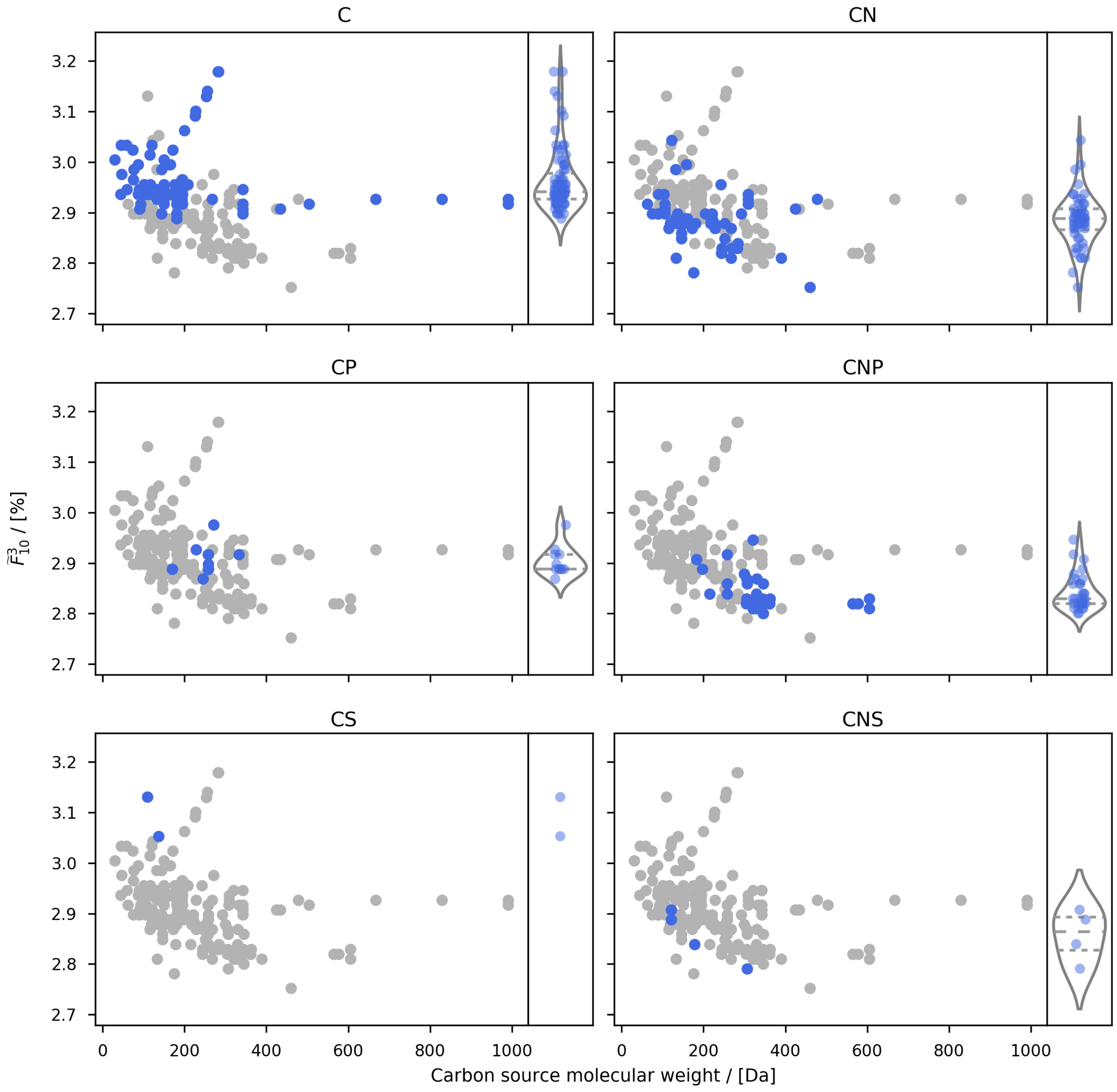


**Supplementary Figure 3**. **PoF vs. carbon source molecular weight grouped by elemental composition.** Each panel highlights the PoF for carbon sources exclusively containing elements as indicated by the panel title plus hydrogen and oxygen. Violin plots show the corresponding distributions.

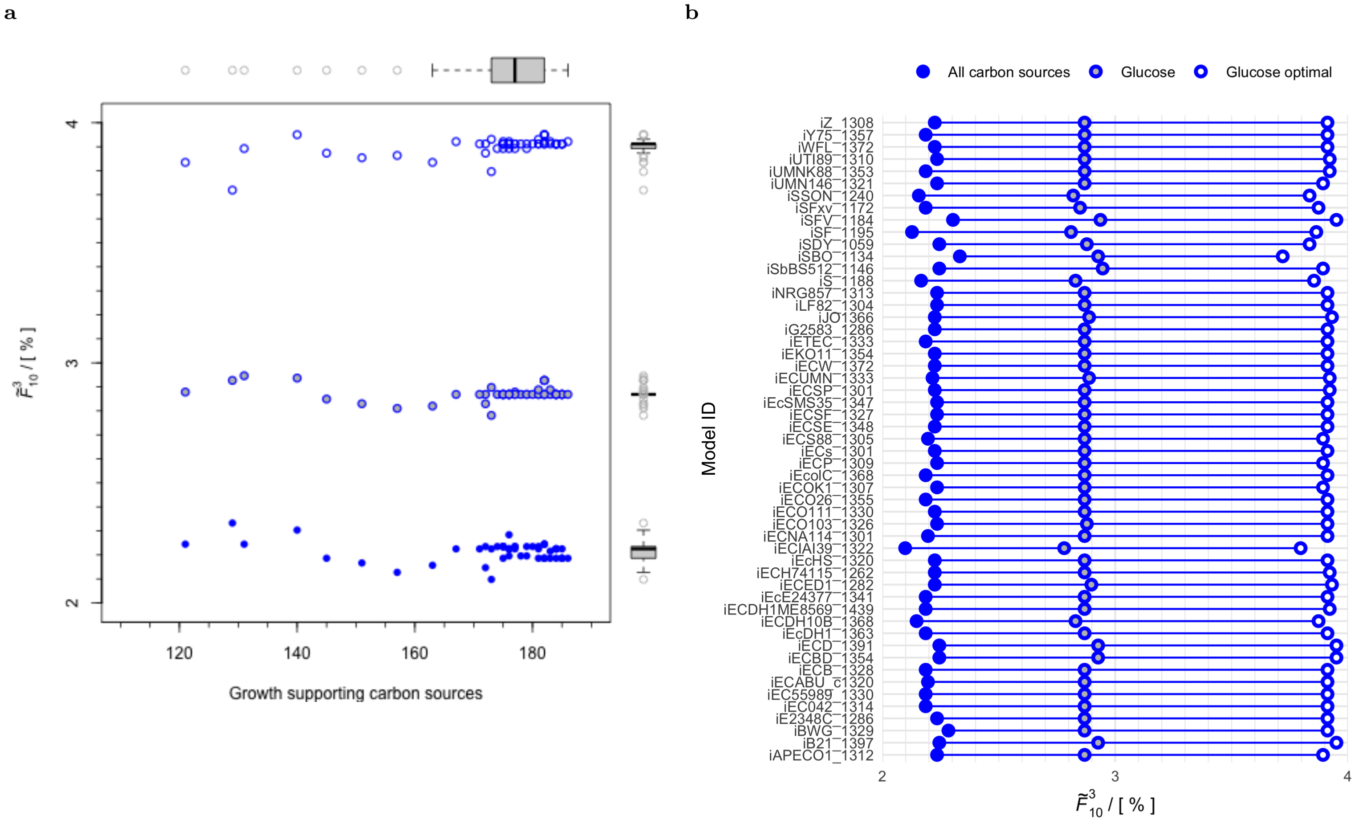


**Supplementary Figure 4**. **PoF and nutritional restriction.** **a**: PoF, as function of the number of growth-supporting environments for multiple GSMMs of *E. coli* and *Shigella* strains. Boxplots on top and to the right summarise the distribution of the data. **b**: PoF for optimal growth on glucose alone, any growth on glucose alone and any growth on all carbon sources present in the respective model. With the exception of *i*SBO_1134 variances among the models are fairly consistent across the three conditions.

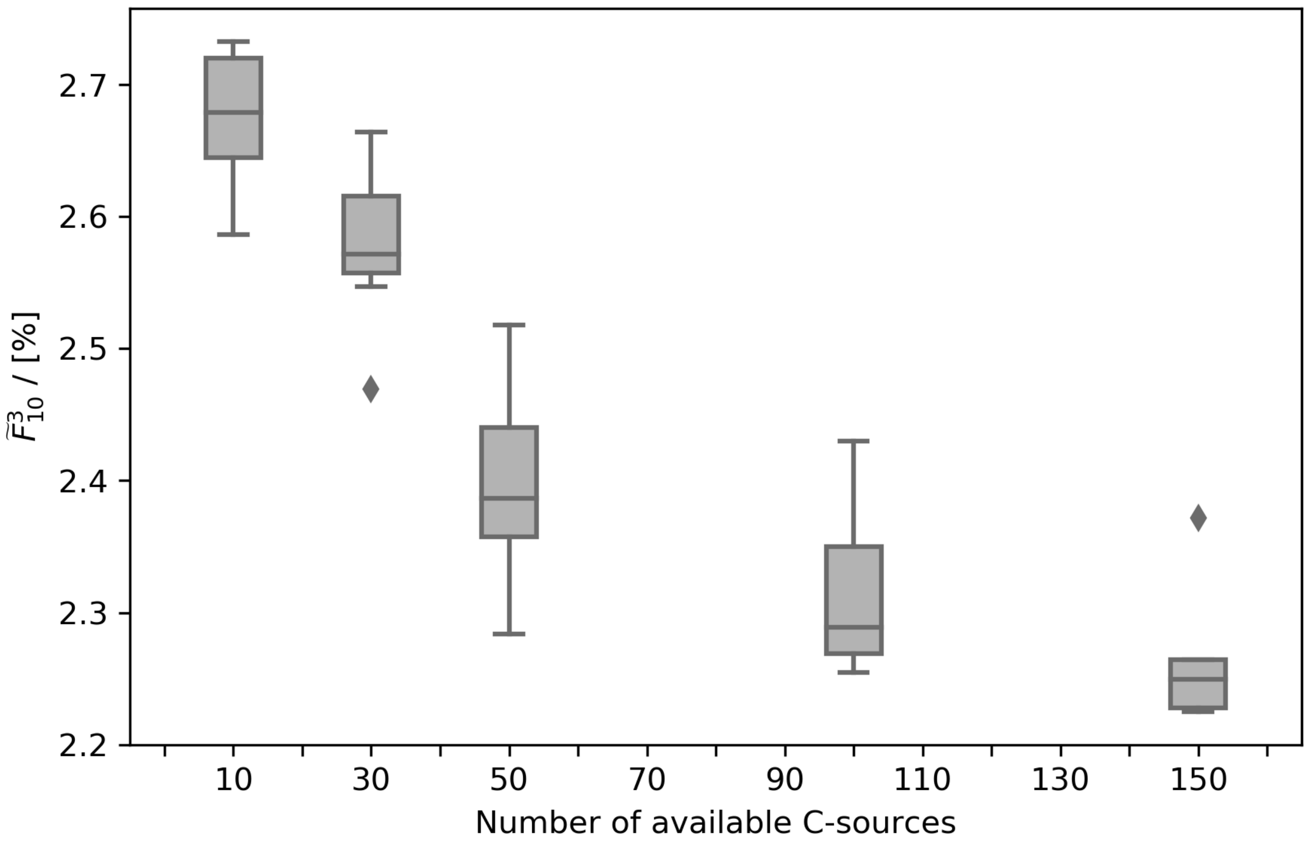


**Supplementary Figure 5**. **PoF as function of the number of utilised carbon sources.** Each box plot illustrates the distribution of the PoF in the GSMM *i*JO1366 across ten runs with randomly selected carbon sources.

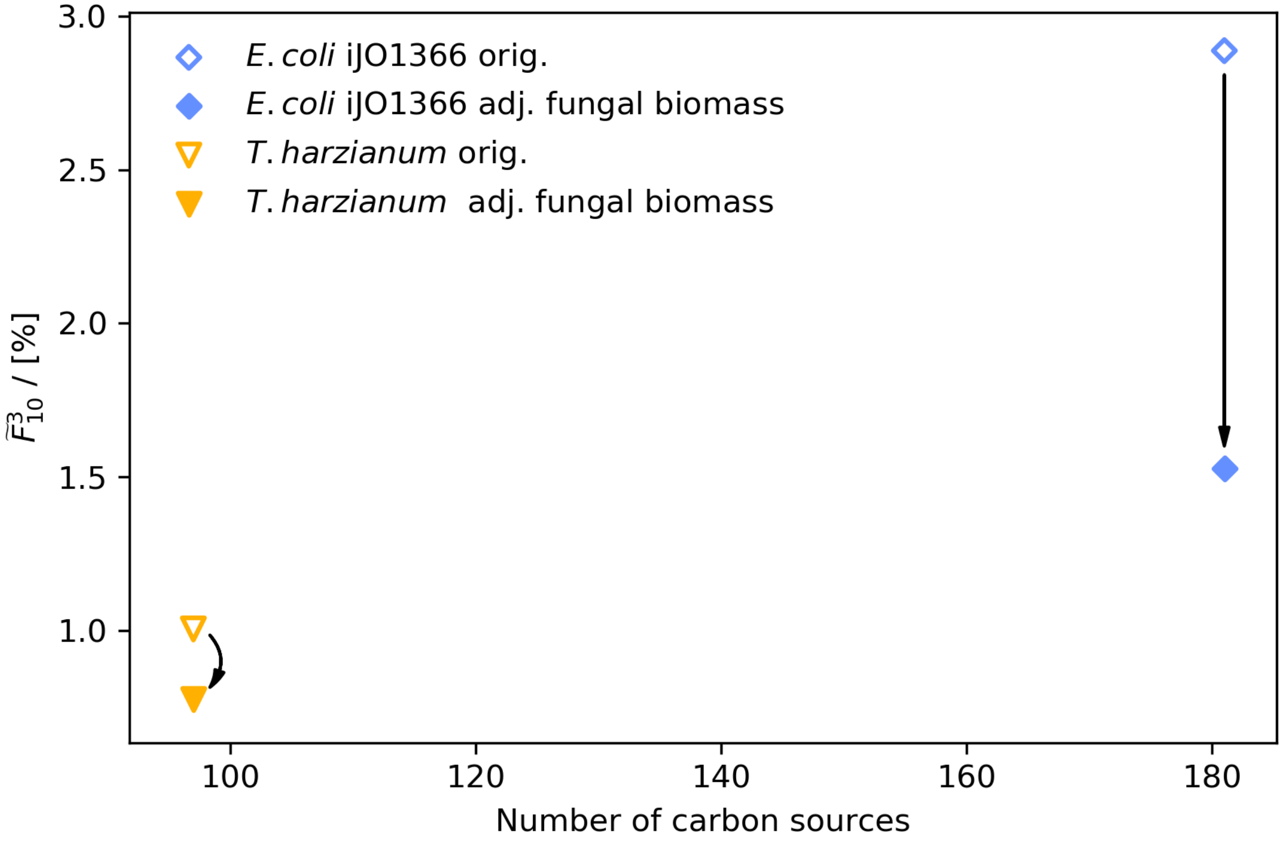


**Supplementary Figure 6**. **Impact of biomass composition on the PoF.** With its growth objective set to the (less detailed – 44 vs. 68 components) biomass reaction used by the fungal models, the *E. coli* GSMM *i*JO1366 showed a drastically decreased PoF. However, it should be noted that four fungal biomass components (ergosterol, zymosterol, chitin, and (1-3)-$\beta$-D-glucan) not present in *i*JO1366 had to be omitted. Removing these from the growth reaction for one of the fungal models (MODEL1604280000, *Trichoderma harzianum*) also reduced its PoF, but to a lesser extent.

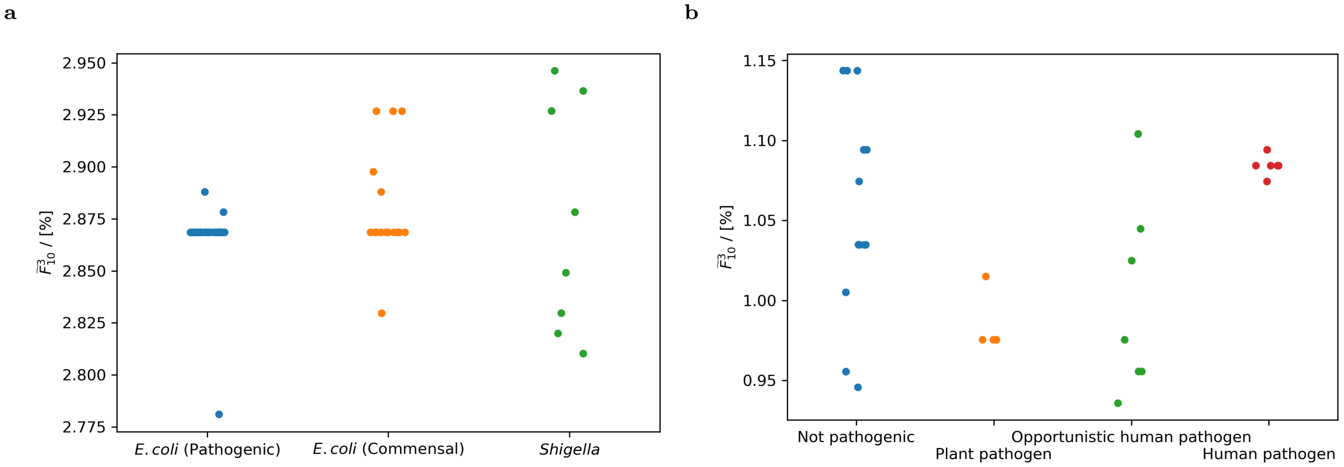


**Supplementary Figure 7**. **PoF of pathogens and non-pathogens.** *Shigella* and pathogenic *E. coli* strains do not exhibit different PoFs compared to non-pathogenic *E. coli* strains **a**. Overall, the fungal species show a similar picture with the variability of the non-pathogens essentially encompassing all other groups **b**. The groups of plant and human pathogens each cluster quite tightly. However, this does not allow to infer general trends given the small number of models and that both groups contained several closely related species (*Fusarium* for plant and *Candida* for human pathogens).

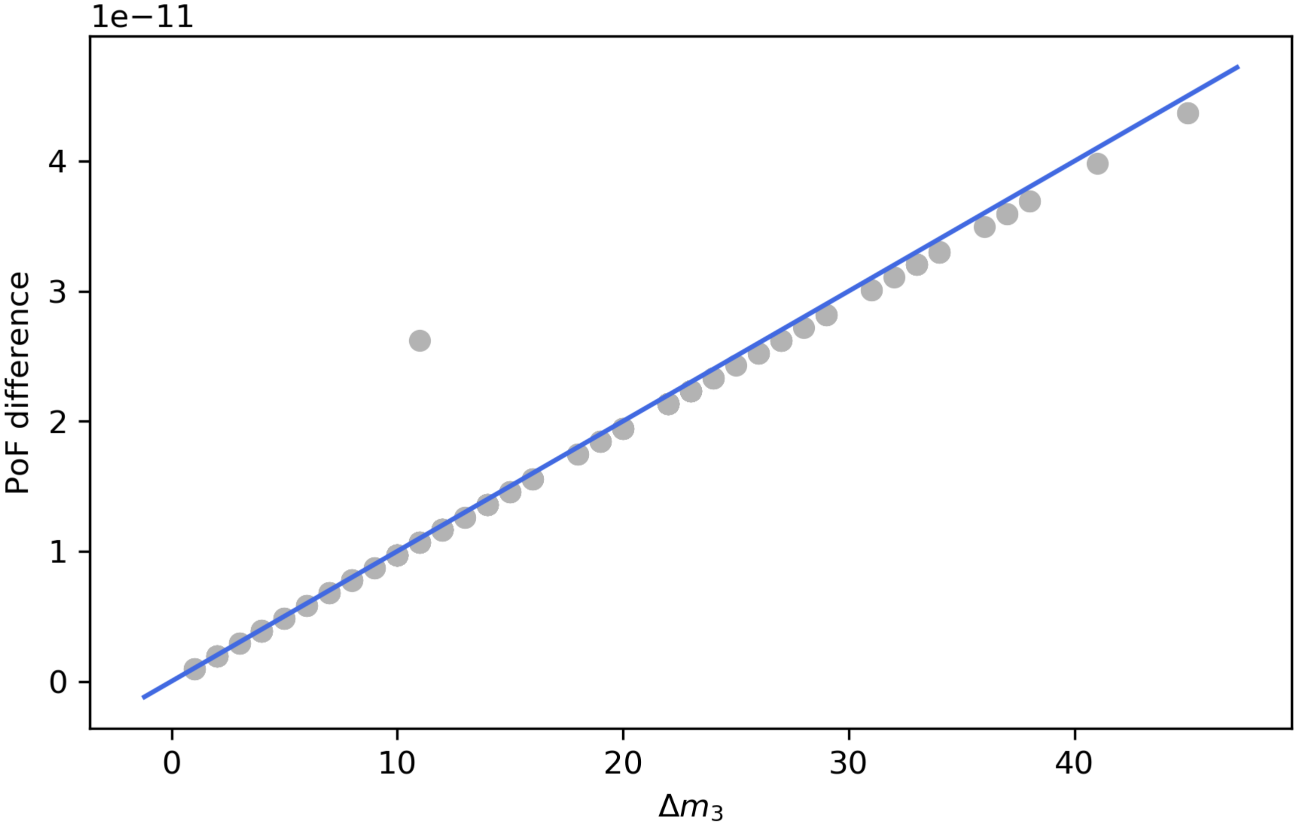


**Supplementary Figure 8**. **Pairwise PoF differences for all *E. coli* pairs with equal $m_1$ and $m_2$.** The blue line is given by eq. (5) with $p = 10^{-12}$.

### SUPPLEMENTARY TABLES

**Supplementary Table 1**. **List of LOF mutations that result in cell death for the toy network in Fig. 1a.** MCSs are highlighted in bold. Note that all elements are super-sets of at least one MCS. Coloured cells indicate the number of possible combinations of LOF mutations. For instance, at $d = 2$ seven out of 15 possible combinations disrupt growth. Finally, the lower part of the table evaluates the PoF, $F$, with the help of eqs. (2) and (3) and using $p = 0.1$.

| $i$ \ $d$ | 1 | 2 | 3 | 4 | 5 | 6 |
|---|---|---|---|---|---|---|
| 1 | $\{\}$ | $\{r_1, r_2\}$ | $\{r_1, r_2, r_3\}$ | $\{r_1, r_2, r_3, r_4\}$ | $\{r_1, r_2, r_3, r_4, r_5\}$ | $\{r_1, r_2, r_3, r_4, r_5, r_6\}$ |
| 2 | | $\{r_4, r_5\}$ | $\{r_1, r_2, r_4\}$ | $\{r_1, r_2, r_3, r_5\}$ | $\{r_1, r_2, r_3, r_4, r_6\}$ | |
| 3 | | $\{r_1, r_6\}$ | $\{r_1, r_2, r_5\}$ | $\{r_1, r_2, r_3, r_6\}$ | $\{r_1, r_2, r_3, r_5, r_6\}$ | |
| 4 | | $\{r_2, r_6\}$ | $\{r_1, r_2, r_6\}$ | $\{r_1, r_2, r_4, r_5\}$ | $\{r_1, r_2, r_4, r_5, r_6\}$ | |
| 5 | | $\{r_3, r_6\}$ | $\{r_1, r_3, r_5\}$ | $\{r_1, r_2, r_4, r_6\}$ | $\{r_1, r_3, r_4, r_5, r_6\}$ | |
| 6 | | $\{r_4, r_6\}$ | $\{r_1, r_3, r_6\}$ | $\{r_1, r_2, r_5, r_6\}$ | $\{r_2, r_3, r_4, r_5, r_6\}$ | |
| 7 | | $\{r_5, r_6\}$ | $\{r_1, r_4, r_6\}$ | $\{r_1, r_3, r_4, r_5\}$ | | |
| 8 | | | $\{r_1, r_5, r_6\}$ | $\{r_1, r_3, r_4, r_6\}$ | | |
| 9 | | | $\{r_2, r_3, r_4\}$ | $\{r_1, r_3, r_5, r_6\}$ | | |
| 10 | | | $\{r_2, r_3, r_6\}$ | $\{r_1, r_4, r_5, r_6\}$ | | |
| 11 | | | $\{r_2, r_4, r_6\}$ | $\{r_2, r_3, r_4, r_5\}$ | | |
| 12 | | | $\{r_2, r_5, r_6\}$ | $\{r_2, r_3, r_4, r_6\}$ | | |
| 13 | | | $\{r_3, r_4, r_6\}$ | $\{r_2, r_3, r_5, r_6\}$ | | |
| 14 | | | $\{r_3, r_5, r_6\}$ | $\{r_2, r_4, r_5, r_6\}$ | | |
| 15 | | | $\{r_4, r_5, r_1\}$ | $\{r_3, r_4, r_5, r_6\}$ | | |
| 16 | | | $\{r_4, r_5, r_2\}$ | | | |
| 17 | | | $\{r_4, r_5, r_3\}$ | | | |
| 18 | | | $\{r_4, r_5, r_6\}$ | | | |
| 19 | | | | | | |
| 20 | | | | | | |
| $f_d$ | 1/6 | 7/15 | 18/20 | 15/15 | 6/6 | 1/1 |
| $w_d$ | 0.354294 | 0.098415 | 0.01458 | 0.001215 | 0.000054 | 0.000001 |
| $\sum_{d'=1}^{d} w_{d'} f_{d'}$ | 0.059049 | 0.104976 | 0.11810 | 0.119313 | 0.119367 | $0.119368 = F$ |

**Supplementary Table 2**. Evaluation of eq. (4) for the toy network in **Fig. 1a**.

| $i$ | $\text{MCS}_i$ | $\mathcal{J}_i$ | $\lvert\mathcal{J}_i\rvert$ | $\mathscr{M}_{\mathcal{J}_i}$ | $\lvert\mathscr{M}_{\mathcal{J}_i}\rvert$ | $(-1)^{\lvert\mathcal{J}_i\rvert-1}p^{\lvert\mathscr{M}_{\mathcal{J}_i}\rvert}$ | $\sum_{i'=1}^{i}(-1)^{\lvert\mathcal{J}_{i'}\rvert-1}p^{\lvert\mathscr{M}_{\mathcal{J}_{i'}}\rvert}$ |
|---|---|---|---|---|---|---|---|
| 1 | $\{r_6\}$ | $\{1\}$ | 1 | $\{r_6\}$ | 1 | $+p$ | $+p$ |
| 2 | $\{r_1, r_2\}$ | $\{2\}$ | 1 | $\{r_1, r_2\}$ | 2 | $+p^2$ | $+p + p^2$ |
| 3 | $\{r_4, r_5\}$ | $\{3\}$ | 1 | $\{r_4, r_5\}$ | 2 | $+p^2$ | $+p + 2p^2$ |
| 4 | $\{r_1, r_3, r_5\}$ | $\{4\}$ | 1 | $\{r_1, r_3, r_5\}$ | 3 | $+p^3$ | $+p + 2p^2 + p^3$ |
| $m = 5$ | $\{r_2, r_3, r_4\}$ | $\{5\}$ | 1 | $\{r_2, r_3.r_4\}$ | 3 | $+p^3$ | $+p + 2p^2 + 2p^3$ |
| 6 | | $\{1, 2\}$ | 2 | $\{r_1, r_2, r_6\}$ | 3 | $-p^3$ | $+p + 2p^2 + p^3$ |
| 7 | | $\{1, 3\}$ | 2 | $\{r_4, r_5, r_6\}$ | 3 | $-p^3$ | $+p + 2p^2$ |
| 8 | | $\{1, 4\}$ | 2 | $\{r_1, r_3, r_5, r_6\}$ | 4 | $-p^4$ | $+p + 2p^2 - p^4$ |
| 9 | | $\{1, 5\}$ | 2 | $\{r_2, r_3, r_4, r_6\}$ | 4 | $-p^4$ | $+p + 2p^2 - 2p^4$ |
| 10 | | $\{2, 3\}$ | 2 | $\{r_1, r_2, r_4, r_5\}$ | 4 | $-p^4$ | $+p + 2p^2 - 3p^4$ |
| 11 | | $\{2, 4\}$ | 2 | $\{r_1, r_2, r_3, r_5\}$ | 4 | $-p^4$ | $+p + 2p^2 - 4p^4$ |
| 12 | | $\{2, 5\}$ | 2 | $\{r_1, r_2, r_3, r_4\}$ | 4 | $-p^4$ | $+p + 2p^2 - 5p^4$ |
| 13 | | $\{3, 4\}$ | 2 | $\{r_1, r_3, r_4, r_5\}$ | 4 | $-p^4$ | $+p + 2p^2 - 6p^4$ |
| 14 | | $\{3, 5\}$ | 2 | $\{r_2, r_3, r_4, r_5\}$ | 4 | $-p^4$ | $+p + 2p^2 - 7p^4$ |
| 15 | | $\{4, 5\}$ | 2 | $\{r_1, r_2, r_3, r_4, r_5\}$ | 5 | $-p^5$ | $+p + 2p^2 - 7p^4 - p^5$ |
| 16 | | $\{1, 2, 3\}$ | 3 | $\{r_1, r_2, r_4, r_5, r_6\}$ | 5 | $+p^5$ | $+p + 2p^2 - 7p^4$ |
| 17 | | $\{1, 2, 4\}$ | 3 | $\{r_1, r_2, r_3, r_5, r_6\}$ | 5 | $+p^5$ | $+p + 2p^2 - 7p^4 + p^5$ |
| 18 | | $\{1, 2, 5\}$ | 3 | $\{r_1, r_2, r_3, r_4, r_6\}$ | 5 | $+p^5$ | $+p + 2p^2 - 7p^4 + 2p^5$ |
| 19 | | $\{1, 3, 4\}$ | 3 | $\{r_1, r_3, r_4, r_5, r_6\}$ | 5 | $+p^5$ | $+p + 2p^2 - 7p^4 + 3p^5$ |
| 20 | | $\{1, 3, 5\}$ | 3 | $\{r_2, r_3, r_4, r_5, r_6\}$ | 5 | $+p^5$ | $+p + 2p^2 - 7p^4 + 4p^5$ |
| 21 | | $\{1, 4, 5\}$ | 3 | $\{r_1, r_2, r_3, r_4, r_5, r_6\}$ | 6 | $+p^6$ | $+p + 2p^2 - 7p^4 + 4p^5 + p^6$ |
| 22 | | $\{2, 3, 4\}$ | 3 | $\{r_1, r_2, r_3, r_4, r_5\}$ | 5 | $+p^5$ | $+p + 2p^2 - 7p^4 + 5p^5 + p^6$ |
| 23 | | $\{2, 3, 5\}$ | 3 | $\{r_1, r_2, r_3, r_4, r_5\}$ | 5 | $+p^5$ | $+p + 2p^2 - 7p^4 + 6p^5 + p^6$ |
| 24 | | $\{2, 4, 5\}$ | 3 | $\{r_1, r_2, r_3, r_4, r_5\}$ | 5 | $+p^5$ | $+p + 2p^2 - 7p^4 + 7p^5 + p^6$ |
| 25 | | $\{3, 4, 5\}$ | 3 | $\{r_1, r_2, r_3, r_4, r_5\}$ | 5 | $+p^5$ | $+p + 2p^2 - 7p^4 + 8p^5 + p^6$ |
| 26 | | $\{1, 2, 3, 4\}$ | 4 | $\{r_1, r_2, r_3, r_4, r_5, r_6\}$ | 6 | $-p^6$ | $+p + 2p^2 - 7p^4 + 8p^5$ |
| 27 | | $\{1, 2, 3, 5\}$ | 4 | $\{r_1, r_2, r_3, r_4, r_5, r_6\}$ | 6 | $-p^6$ | $+p + 2p^2 - 7p^4 + 8p^5 - p^6$ |
| 28 | | $\{1, 2, 4, 5\}$ | 4 | $\{r_1, r_2, r_3, r_4, r_5, r_6\}$ | 6 | $-p^6$ | $+p + 2p^2 - 7p^4 + 8p^5 - 2p^6$ |
| 29 | | $\{1, 3, 4, 5\}$ | 4 | $\{r_1, r_2, r_3, r_4, r_5, r_6\}$ | 6 | $-p^6$ | $+p + 2p^2 - 7p^4 + 8p^5 - 3p^6$ |
| 30 | | $\{2, 3, 4, 5\}$ | 4 | $\{r_1, r_2, r_3, r_4, r_5\}$ | 5 | $-p^5$ | $+p + 2p^2 - 7p^4 + 7p^5 - 3p^6$ |
| $2^m - 1 = 31$ | | $\{1, 2, 3, 4, 5\}$ | 5 | $\{r_1, r_2, r_3, r_4, r_5, r_6\}$ | 6 | $+p^6$ | $+p + 2p^2 - 7p^4 + 7p^5 - 2p^6 = F$ |

**Supplementary Table 3**. Fungal BioModels [1] accession IDs

| Accession ID | Species |
|---|---|
| MODEL1604280000 | Trichoderma harzianum |
| MODEL1604280001 | Penicillium chrysogenum |
| MODEL1604280003 | Fusarium verticillioides |
| MODEL1604280004 | Trichoderma citrinoviride |
| MODEL1604280005 | Chaetomium globosum |
| MODEL1604280006 | Candida tropicalis |
| MODEL1604280007 | Pichia guilliermondii |
| MODEL1604280010 | Trichoderma longibrachiatum |
| MODEL1604280012 | Aspergillus oryzae |
| MODEL1604280013 | Magnaporthe grisea |
| MODEL1604280016 | Aspergillus clavatus |
| MODEL1604280017 | Yarrowia lipolytica |
| MODEL1604280018 | Fusarium oxysporum |
| MODEL1604280019 | Aspergillus terreus |
| MODEL1604280022 | Trichoderma asperellum |
| MODEL1604280024 | Trichoderma reesei |
| MODEL1604280025 | Nectria haematococca |
| MODEL1604280026 | Schizosaccharomyces japonicus |
| MODEL1604280028 | Debaryomyces hansenii |
| MODEL1604280029 | Aspergillus fumigatus |
| MODEL1604280031 | Fusarium graminearum |
| MODEL1604280033 | Candida glabrata |
| MODEL1604280038 | Trichoderma virens |
| MODEL1604280039 | Lodderomyces elongisporus |
| MODEL1604280041 | Schizosaccharomyces pombe |
| MODEL1604280043 | Candida lusitaniae |
| MODEL1604280046 | Neurospora crassa |
| MODEL1604280049 | Pichia stipitis |
| MODEL1604280052 | Candida albicans |
| MODEL1604280055 | Pichia pastoris |

**Supplementary Table 4**. Minimal media.

| Medium component | Reaction ID | *E. coli*, *Shigella*, *Salmonella* | fungi |
|---|---|---|---|
| Calcium | EX_ca2_e | + | - |
| Chloride | EX_cl_e | + | - |
| $Co^{2+}$ | EX_cobalt2_e | + | - |
| $Cu^{2+}$ | EX_cu2_e | + | - |
| $Fe^{2+}$ | EX_fe2_e | + | - |
| D-Glucose | EX_glc__D_e | + | + |
| $H_2O$ | EX_h2o_e | + | - |
| $H^+$ | EX_h_e | + | - |
| $K^+$ | EX_k_e | + | - |
| Mg | EX_mg2_e | + | - |
| $Mn^{2+}$ | EX_mn2_e | + | - |
| Molybdate | EX_mobd_e | + | - |
| Ammonium | EX_nh4_e | + | + |
| $Ni^{2+}$ | EX_ni2_e | + | - |
| $O_2$ | EX_o2_e | + | + |
| Phosphate | EX_pi_e | + | + |
| Sulfate | EX_so4_e | + | + |
| Zinc | EX_zn2_e | + | - |

**Supplementary Table 5**. Auxotrophic *E. coli* and *Shigella* strains and the respective supplements required for growth.

| Strain | Model ID | Supplement | Exchange reaction ID |
|---|---|---|---|
| Shigella boydii Sb227 | iSBO_1134 | Thiamin, Niacin | EX_thm_e, EX_nac_e |
| Shigella flexneri 2002017 | iSFxv_1172 | Niacin | EX_nac_e |
| Shigella sonnei Ss046 | iSSON_1240 | Niacin | EX_nac_e |
| Shigella flexneri 5 str. 8401 | iSFV_1184 | Niacin | EX_nac_e |
| Shigella flexneri 2a str. 301 | iSF_1195 | Methionine, Niacin | EX_met__L_e, EX_nac_e |
| Escherichia coli UMN026 | iECUMN_1333 | Niacin | EX_nac_e |
| Shigella boydii CDC 3083-94 | iSbBS512_1146 | Thiamin | EX_thm_e |
| Shigella flexneri 2a str. 2457T | iS_1188 | Niacin | EX_nac_e |
| Escherichia coli str. K-12 substr. DH10B | iECDH10B_1368 | Leucin | EX_leu__L_e |

**Supplementary Table 6**. Auxotrophic *Salmonella* strains and the respective supplements required for growth. NMN, Nicotinamide mononucleotide

| Strain | Model ID | Supplement | Exchange reaction ID |
|---|---|---|---|
| EC20120219 | YS__Enteritidis__EC20120219 | Tryptophan | Auxotrophy_trp__L_e |
| ATCC_51960 | YS__Albany__ATCC_51960 | Xanthine | Auxotrophy_xan_e |
| EC20121750 | YS__Enteritidis__EC20121750 | Histidine | Auxotrophy_his__L_e |
| 99_7863 | YS__Paratyphi A__99_7863 | NMN | Auxotrophy_nmn_e |
| EC20120597 | YS__Enteritidis__EC20120597 | Tryptophan | Auxotrophy_trp__L_e |
| SA20090877 | YS__Enteritidis__SA20090877 | Tryptophan | Auxotrophy_trp__L_e |
| EC20120229 | YS__Enteritidis__EC20120229 | Tryptophan | Auxotrophy_trp__L_e |
| EC20121753 | YS__Enteritidis__EC20121753 | Tryptophan | Auxotrophy_trp__L_e |
| [NA] | YS__Paratyphi A__9_65 | NMN | Auxotrophy_nmn_e |
| 01_1852 | YS__Paratyphi A__01_1852 | NMN | Auxotrophy_nmn_e |
| [NA] | YS__Paratyphi A__98_9652 | NMN | Auxotrophy_nmn_e |
| SA20094521 | YS__Enteritidis__SA20094521 | Tryptophan | Auxotrophy_trp__L_e |
| EC20120686 | YS__Enteritidis__EC20120686 | Tryptophan | Auxotrophy_trp__L_e |
| EC20130348 | YS__Enteritidis__EC20130348 | Tryptophan | Auxotrophy_trp__L_e |
| U288 | YS__Typhimurium__U288 | Histidine | Auxotrophy_his__L_e |
| EC20121748 | YS__Enteritidis__EC20121748 | Tryptophan | Auxotrophy_trp__L_e |
| A103(ParaA) | YS__Paratyphi A__A103(ParaA) | NMN | Auxotrophy_nmn_e |
| SA19942384 | YS__Enteritidis__SA19942384 | Tryptophan | Auxotrophy_trp__L_e |
| EC20122031 | YS__Enteritidis__EC20122031 | Tryptophan | Auxotrophy_trp__L_e |
| SA20093421 | YS__Enteritidis__SA20093421 | Tryptophan | Auxotrophy_trp__L_e |
| SA20090435 | YS__Enteritidis__SA20090435 | Tryptophan | Auxotrophy_trp__L_e |
| A61_149 | YS__Paratyphi A__A61_149 | NMN | Auxotrophy_nmn_e |
| EC20090195 | YS__Enteritidis__EC20090195 | Tryptophan | Auxotrophy_trp__L_e |
| SA19930684 | YS__Enteritidis__SA19930684 | Tryptophan | Auxotrophy_trp__L_e |
| 138_69 | YS__Paratyphi A__138_69 | NMN | Auxotrophy_nmn_e |

*
†

[1] R. S. Malik-Sheriff, M. Glont, T. V. N. Nguyen, K. Tiwari, M. G. Roberts, A. Xavier, M. T. Vu, J. Men, M. Maire, S. Kananathan, E. L. Fairbanks, J. P. Meyer, C. Arankalle, T. M. Varusai, V. Knight-Schrijver, L. Li, C. Dueñas-Roca, G. Dass, S. M. Keating, Y. M. Park, N. Buso, N. Rodriguez, M. Hucka, and H. Hermjakob, Nucleic Acids Research 10.1093/nar/gkz1055 (2019).
[2] M. P. Gerstl, S. Klamt, C. Jungreuthmayer, and J. Zanghellini, Bioinformatics **32**, 730 (2016).
[3] J. D. Orth, T. M. Conrad, J. Na, J. A. Lerman, H. Nam, A. M. Feist, and B. Ø. Palsson, Mol Syst Biol **7**, 535 (2011).
[4] J. D. Orth, R. M. T. Fleming, and B. O. Palsson, EcoSal Plus **4**, 10.1128/ecosalplus.10.2.1 (2010).
